# Strain dependent structural effects and *in vivo* efficacy of enterovirus-D68 inhibitors

**DOI:** 10.1101/2021.12.24.474126

**Authors:** Thomas Lane, Jianing Fu, Barbara Sherry, Bart Tarbet, Brett L. Hurst, Olga Riabova, Elena Kazakova, Anna Egorova, Penny Clarke, J. Smith Leser, Joshua Frost, Michael Rudy, Ken Tyler, Thomas Klose, Richard J. Kuhn, Vadim Makarov, Sean Ekins

## Abstract

Acute flaccid myelitis (AFM) leads to loss of limb control in young children and is likely due to Enterovirus-D68 (EV-D68), for which there is no current treatment. We have developed a lead isoxazole-3-carboxamide analog of pleconaril (11526092) which displayed potent inhibition of the pleconaril-resistant CVB3-Woodruff (IC_50_ 6-20 nM), EV-D68 (IC_50_ 58 nM), and other enteroviruses. A mouse respiratory model of EV-D68 infection, in which pleconaril is inactive, showed decreased viremia of 3 log units as well as statistically significant 1 log reduction in lung titer reduction at day 5 after treatment with 11526092. A cryo-electron microscopy (cryo-EM) structure of EV-D68 in complex with 11526092 suggests that the increased potency may be due to additional hydrophobic interactions. Cryo-EM structures of 11526092 and pleconaril demonstrate destabilization of EV-D68 (MO strain) compared to the previously described stabilization of EV-D68 (Fermon strain) with pleconaril, illustrating clear strain dependent mechanisms of this molecule. 11526092 represents a more potent inhibitor *in vitro* with *in vivo* efficacy providing a potential future treatment for EV-D68 and AFM, suggesting an improvement over pleconaril for further optimization.

**One-Sentence Summary:** 11526092 demonstrates protein destabilization, improved *in vitro* potency and *in vivo* efficacy when compared with pleconaril against EV-D68.

---

Enteroviruses (EVs), coxsackieviruses (CVs) and rhinoviruses (RVs) of the *Enterovirus* genus of the *Picornaviridae* family frequently cause lower respiratory infections (*1–4*) which represent the main cause of death in low-income countries and the third largest cause of death worldwide. EVs and CVs also cause a wide range of acute and chronic diseases, such as aseptic meningitis, encephalitis, hand-foot-and-mouth disease, conjunctivitis, diarrhea, herpetic angina, acute and chronic myocarditis, etc. (*5*). Enterovirus D68 (EV-D68) shares important biological and molecular properties with other EVs and RVs and has emerged as an important global public health threat (*6*). EV-D68, like all members of the *Enterovirus* genus, is a non-enveloped, single-stranded, positive sense RNA virus of the family Picornaviridae (*7*). Despite identification in 1962, disease outbreaks were not widely reported until the early 2000s. Beginning with an outbreak of severe acute respiratory disease in the Philippines in 2008–2009, an increase in confirmed cases of EV-D68 infection in children (mean age of onset between 3–8 years old (*8*)) led to the classification of EV-D68 as a reemerging pathogen (*9*). 1,395 cases of EV-D68 respiratory infections were confirmed between August 2014-January 2015 by the CDC (*10*). Over 1000 additional cases of EV-D68 respiratory disease were also confirmed in Canada, Europe, Asia, and South America (*11*). Accumulating evidence supports an association between EV-D68 and Acute flaccid myelitis (AFM) (*12*), a disease which afflicts approximately 1% of EV-D68 patients and which is similar to poliomyelitis (*13*). The majority (80–90%) of children with AFM also experience viral prodromal symptoms of fever and upper respiratory illness in the week prior to the onset of limb weakness (*8*). The severity of respiratory disease, however, does not appear to correlate with the development of paralysis (*12*). Other viruses that have been associated with outbreaks of acute flaccid weakness and AFM include enterovirus A71 (EV-A71), West Nile virus, Japanese encephalitis virus, and the wild-type poliovirus (*14–16*). EV-D68 appears to occur in a cyclic pattern with a 2-year interval (*8, 12, 17*). Clearly more research is needed (*18*), as are treatments to decrease morbidity and mortality resulting from these infections. Current treatment of EV, CV and RV infections aims to reduce and shorten symptoms (e.g. fever and pain, fatigue, and nasal blockage in the case of common cold). Although several molecules have been evaluated and have shown promise *in vitro*, they have all failed in *in vivo* models. For example, fluoxetine, while effective in inhibiting EV-D68 replication *in vitro* did not reduce the viral load in the mouse model (virus infection in 2-day old pups) of EV-D68–associated AFM or improve motor function, and ultimately failed in clinical studies (*19, 20*). The most promising results against EVs have been obtained with the capsid-binding inhibitors such as pleconaril and vapendavir (*21, 22*). However, due to insufficient effectiveness and/or side effects in the clinic, neither of these inhibitors has been approved by the FDA (*23*). Pleconaril binds in a hydrophobic pocket within VP1 of EVs and RVs, stabilizes the viral capsid and prevents viral adsorption and/or uncoating (*24*). The pleconaril-resistant strain of coxsackievirus B3 contains two genetic changes in VP1 at position 92 from Ile to Leu or Met and at position 207 from Ile to Val (*21, 25*). The mutation at position 192 of VP1 renders EV-71 resistant to pyridyl imidazolidinone, indicating position 192 of VP1 plays an important role in interacting with this antiviral compound (*24, 26*). Therefore there is no approved antiviral treatment for EVs, CVs and RVs, and with the exception of the poliovirus vaccine no prophylaxis, to prevent the millions of lost school and working days (*27*). We have recently developed novel isoxazole-3-carboxamide analogs of pleconaril which we described with activity against CVs and RVs, one of which, 3-(3-Methyl-4-(3-(3-N,N-dimethylcarbamoyl-isoxazol-5-yl)propoxy)phenyl)-5-trifluoromethyl-1,2,4-oxadiazole (11526092), possessed acceptable pharmacokinetics in mice (*28*) and which we now describe as displaying potent inhibition across various EVs *in vitro*. We have now shown using cryo-electron microscopy (cryo-EM) that pleconaril binds differently to the Fermon and MO strains of EV-D68. In addition, 11526092 binds to the EV-D68 VP1 from the MO strain similarly to pleconaril and has demonstrated promising antiviral activity against EV-D68 in a mouse respiratory model of EV-D68 infection for AFM.

## Results

Following our earlier studies in CVs and RVs (*28*) we focused on 3 isoxazole-3-carboxamide derivatives of pleconaril (Fig. 1). These molecules had lower predicted physicochemical properties such as logP, logD and pKa than pleconaril but increased molecular weight (Table S1). When these new molecules were tested *in vitro* using CVB3-Woodruff (Supplementary Materials and Methods), 11526091 and 11526092 IC_50_ values were statistically significantly lower than for pleconaril (p<0.0001) and this was independent of adding drug pre- or post-infection (Table 1). Interestingly, 11526092 also displayed the greatest antiviral effect in preliminary work with human rhabdomyosarcoma RD cells infected with EV-D68 (Fig. S1). 11526092 was then profiled against several viruses and showed activity against CVB3, EV-D68, polio, less activity against EV-71 and influenza H1N1, and no activity against the recently identified SARS-CoV-2 in Vero cells (Table 2). The expected target of 11526092 is the capsid protein VP1, which is structurally highly similar in multiple proteins encoded by the *Picornaviridae* family (Fig. S2). The potency *in vitro* against EV-D68 and maEV-D68 (EC_50_ 23-60 nM, Table 2) prompted us to pursue 11526092 further. The ADME properties for 11526092 were determined (Table 3) and it demonstrated low solubility, inhibition of CYP2C9 and CYP2C19 and poor mouse microsomal stability. In human liver microsomes 11526092 is stable while the compound is highly protein bound. The Caco-2 data suggest that 11526092 is likely a P-gp substrate and this may affect the pharmacokinetics of 11526092. Neither appreciable hERG inhibition or PXR agonist activity were observed (Table 3). The maximum tolerated dose for 11526092 was determined in 2-week-old and 10-day-old AG129 mice dosed either by oral (PO) or intraperitoneal (IP) administration, respectively (Fig. S3). The maximum tolerated dose of 11526092 in AG129 mice was found to be >100 mg/kg when given PO and >10 mg/kg when given via IP. This study was followed by determining the efficacy of 11526092 for treatment of an EV-D68 respiratory infection in four-week-old AG129 mice. The EV-D68 respiratory model (*29*), based on intranasal infection of 4-week-old AG129 mice, is a non-lethal model that includes viremia with rapid tissue distribution of virus, including high virus titers in lung tissues and an elevation of proinflammatory cytokines in the lung. Treatment with 11526092 at 100 mg/kg/d alone provided significant protection from weight loss (as a percentage of initial body weight) following infection (Fig. S4). Titers of virus in the blood from mice following 11526092 treatment and virus challenge showed all doses were able to reduce viremia on days 1 and 3 post-infection (p.i.) (Fig. 2A). No virus was detected for any group on day 5 p.i. Lung titers on days 1, 3, and 5 post-infection (p.i.) were measured after treatment and these results showed that only the guanidine-treated group showed a reduction in virus titers for days 1 and 3 p.i. compared to placebo (Fig. 2B). However, on day 5 p.i. mice treated with the 30 mg dose of 11526092 also showed a significant reduction in virus titer (Fig. 2B). Interestingly, during animal model development (*29*) pleconaril did not demonstrate any antiviral activity. Early activation of the innate immune system is essential for immediate control of a respiratory virus replication and spread. Production of antiviral type I interferons (IFNs), for example, by lung epithelial cells, macrophages, and dendritic cells leads to the expression of IFN-stimulated genes with antiviral and immune-modulatory functions (*30, 31*). All doses of 11526092 significantly reduced lung concentrations of IL-3, IL-5, GM-CSF, and RANTES on day 1 after treatment and challenge infection. Additional reductions in cytokine concentrations after treatment and infection included IFN-γ and MCP-1 on day 1 for the 10 and 100 mg/kg treatment groups, respectively. An increase in IL-1α, IL-1β, IL-4, and IL-6 were observed on day 3 for the 100 mg/kg treatment groups (Fig. S5 and Supplementary text).

**Fig. 1.**
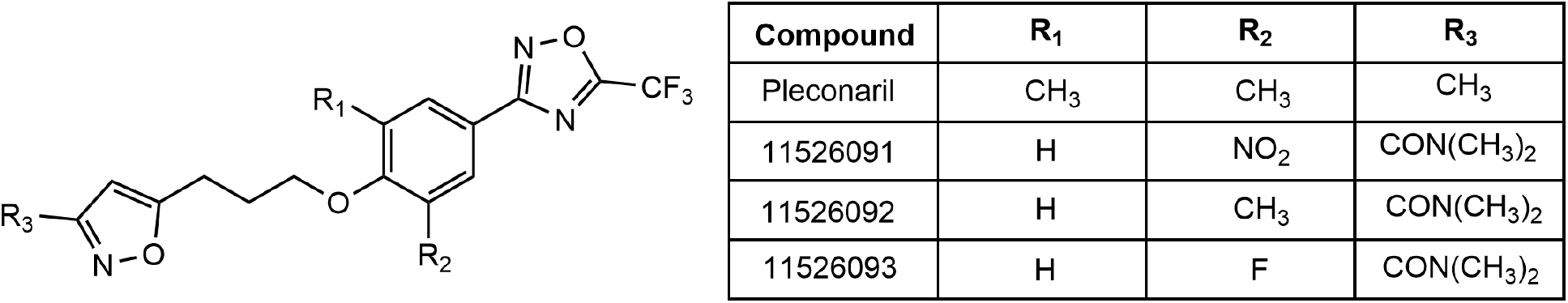
Structures of isoxazole-3-carboxamide pleconaril derivatives.

**Fig. 2.**
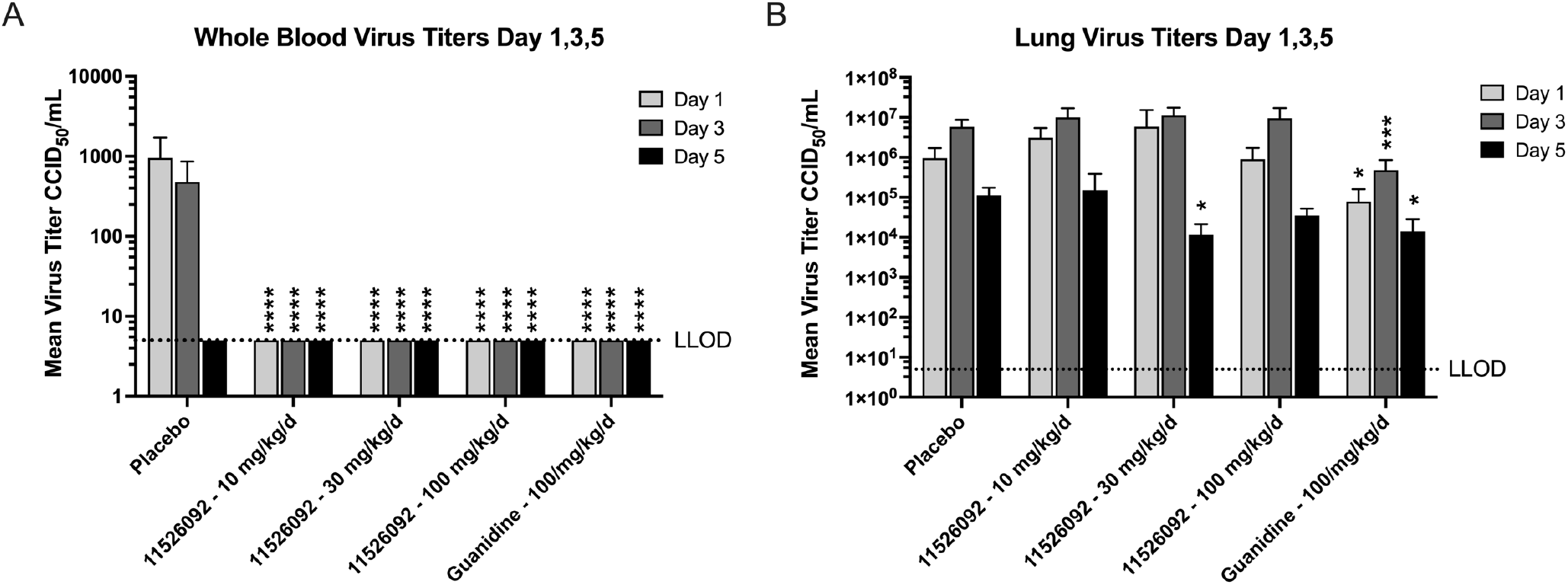
AFM respiratory model viremia levels in A. whole blood and B. lung. Infectious virus was determined for viremia and viral lung titers by assessing the infection of human rhabdomyosarcoma (RD) cells in microtiter plates. Fifty percent cell culture infectious doses (CCID50) were converted to CCID50 per gram of lung prior to statistical analysis. Ordinary ANOVA followed by a Dunnett’s T3 multiple comparisons test was performed in Graphpad Prism 9.2 for macOS. Asterisks represent statistical significance using the GP style, where P values of ≤0.0001, ≤0.001, ≤0.01, ≤0.05 and ≥0.05 are summarized with “****”, “***”, “**”, “*” and ns, respectively. Error bars represent the SEM.

**Table 1.** Isoxazole-3-Carboxamide derivatives testing with CVB3-Woodruff versus pleconaril. “±” represents the SEM (n ≥ 6) as calculated in Prism 9.2. As not all the compounds reached total inhibition at the concentrations tested, top and bottom were constrained to be shared for each MOI tested. Values were statistically significantly lower than for pleconaril (p<0.0001) for 11526091 and 11526092, independent of adding drug pre- or post-infection.

| Name | Inhibitory concentration $IC_{50}$ against CVB3-Woodruff, $\mu M$ | | |
| --- | --- | --- | --- |
|  | Adding drug to virus and then infecting at indicated multiplicity of infection (MOI): |  | Infecting at low MOI and adding drug after infection is established: |
|  | 1 PFU / cell | 3 PFU / cell | 0.1 PFU / cell |
| **Pleconaril** | $0.048 \pm 0.004$ | $0.187 \pm 0.015$ | $0.156 \pm 0.009$ |
| **11526091** | $0.006 \pm 0.000$ | $0.010 \pm 0.001$ | $0.016 \pm 0.001$ |
| **11526092** | $0.006 \pm 0.000$ | $0.010 \pm 0.001$ | $0.016 \pm 0.041$ |
| **11526093** | $0.049 \pm 0.008$ | $0.055 \pm 0.009$ | $0.450 \pm 0.041$ |

**Table 2.** NIAID testing of 11526092 against several viruses of interest (± SD when replicated, N.D. = not determined)

| **Virus Serotype** | **Strain Name** | **50%**<br>**Effective**<br>**concentration**<br>**EC<sub>50</sub> (μM)** | **90%**<br>**Effective**<br>**concentration**<br>**EC<sub>90</sub> (μM)** | **50%**<br>**Cytotoxic**<br>**concentration**<br>**CC<sub>50</sub> (μM)** |
| --- | --- | --- | --- | --- |
| Coxsackie B3 | HA 201933 | 0.92 ± 0.53 | N.D. | 7.38 ± 0.72 |
| Coxsackie B3 | Nancy | 1.27 | N.D. | 8.0 |
| EV-D68 | US/KY/14-18953 | 0.059 ± 0.023 | 0.078 | 6.6 ± 1.33 |
| ma-EV-D68 | US/MO/14-18949 (Mouse adapted) | 0.028 | N.D. | 9.4 |
| EV-71 | Tainan/4643/98 | 2.21±2.87 | N.D. | 4.37 ± 4.48 |
| Polio virus 1 | Mahoney | 0.48±0.44 | 2.83 | 9.96 ± 2.78 |
| Influenza H1N1 | California/07/2009 | N.D. | 10.37 | 44.77 |
| SARS-CoV-2 | USA/WA1/2020 | >7.54 | N.D. | 7.54 |

**Table 3.** ADME properties for 11526092. * CYP3A4 demonstrates activation with 11526092.

| ADME property | Data |
| --- | --- |
| **Solubility** | 0.11 $\mu\text{M}$ in Buffer pH 7.4, 12.05 $\mu\text{M}$ in Fasted State Simulated Intestinal Fluid (FaSSIF), 0.55 $\mu\text{M}$ in Fasted State Simulated Gastric Fluid (FaSSGF). |
| **CYP inhibition** | CYPs1A2, 2D6, 3A4* ($>50 \mu\text{M}$), 2C9 (5 $\mu\text{M}$), 2C19 (2.1 $\mu\text{M}$) |
| **Mouse liver microsomes** | $t_{1/2}$ = 19.6 min (unstable),<br>$\text{CL}_{\text{int}}$ = 70.6 $\mu\text{L}/\text{min}/\text{mg}$ protein |
| **Human liver microsomes** | $T_{1/2}$ = 55.3 min (stable), 25.1 $\mu\text{L}/\text{min}/\text{mg}$ protein |
| **Mouse plasma protein binding** | 99.8% bound, 88.1% stability |
| **Human plasma protein binding** | 99.7% bound, 93.9% stability |
| **Mouse plasma stability** | 81.1% @ 5 hr, 37°C |
| **Human plasma stability** | 93.9% @ 5 hr, 37°C |
| **Caco-2** | Papp A-B = $8.8 \times 10^{-4} \text{ cm/s}$; B-A = $30.9 \times 10^{-4} \text{ cm/s}$ Efflux ratio = 3.5 (likely P-gp substrate) |
| **hERG inhibition** | $\text{IC}_{50} > 100 \mu\text{M}$ |
| **PXR** | $\text{EC}_{50} > 100 \mu\text{M}$ |

The antiviral efficacy of 11526092 for treatment of mouse-adapted (ma) EV-D68 neurological infection in 10-day-old AG129 mice was assessed in two studies where 11526092 was administered either 2 hours pre-infection or 4 hours post-infection (Fig. S6). None of the 1526092-treated mice were protected from challenge infection in this model. Guanidine, used as positive control, protected >80% of the mice in the post-challenge study. None of the treatment groups provided significant protection from weight loss (Fig. S7) and none of the 1526092-treated mice showed a significant reduction in neurological scores while guanidine reduced the neurological score on day 3 post-infection. Only the guanidine treated group survived past day 4 in the post challenge study. None of the 1526092-treated mice showed a significant reduction in virus titers on days 1 or 3. Only the guanidine treated mice survived to day 5 post-infection. None of the prechallenge 1526092-treated mice were protected from challenge infection. Guanidine protected 100% of the mice and provided a significant protection from weight loss with pretreatment (Fig. S7). Treatment with 11526092 increased the mean day of death for mice that died following challenge, although that increase in survival was not significant. The mean day of death for the placebo group was 5.5 days and the mean day of death for each treatment group was 6.3, 5.4, and 4.3 days, for the 1, 3 and 10 mg 11526092-treated groups, respectively. Too few animals in the placebo and 11526092-treated groups survived past day 5 to complete statistical comparisons for later time points.

We have also investigated the cryo-EM structure of EV-D68 MO in complex with 11526092 in order to demonstrate the mechanism of inhibition. The capsid of EV-D68 consists of 60 copies of the viral protein (VP) 1, VP2, VP3 and VP4 and has an icosahedral pseudo-T=3 symmetry (*32–34*). VP 1, 2 and 3 follow the typical jelly roll-fold pattern and form the capsid surface, whereas

VP4 is located inside the capsid and contributes to the stability of the capsid and genome (*35–38*). Each of the five-fold axes is surrounded by a deep depression called the “canyon” which is the receptor binding site for a large number of EV’s (*33-35, 39*). Underneath the canyon and within VP1 there is a hydrophobic region called the pocket (*32, 35, 39*). 11526092 binds to the capsid VP1 (Fig. 3) in a manner like pleconaril and this is also similar to the initial docking prediction made with previous cryo-EM structures (Fig. S8) (*32*). We have also solved the structure of EV-D68 and another isoxazole-3-carboxamide analog of pleconaril, namely 11526093, although the density of compound is too weak to fit into the pocket (Supplemental text).

**Fig. 3.**
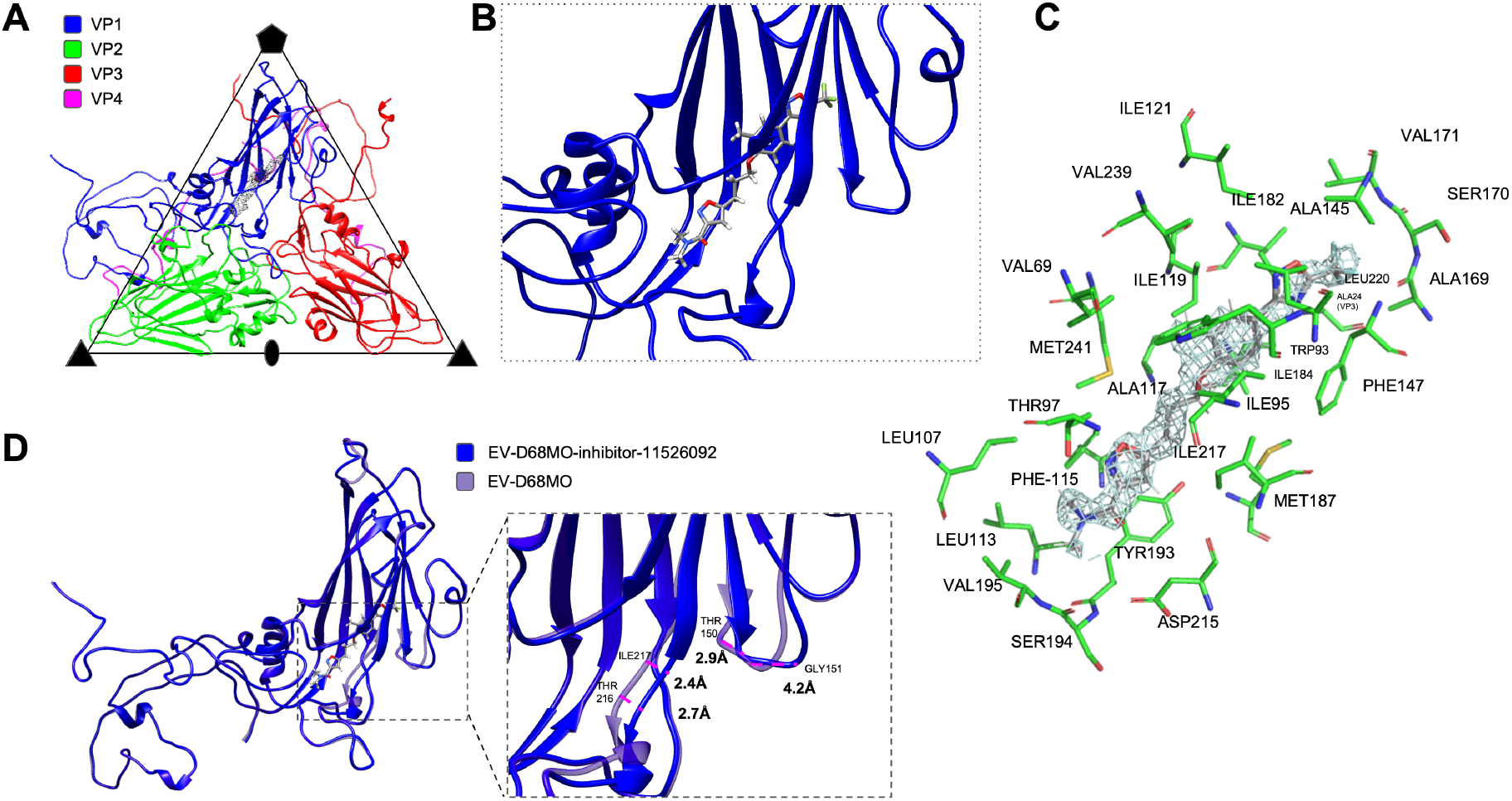
Cryo-EM structure of Enterovirus 68 (EV-D68) inhibitor 11526092 bound to the capsid Viral Protein 1 (VP1) of EV-D68 (MO strain). (A) An icosahedral asymmetric unit of the EV-D68-inhibitor 11526092 complex. VP1, VP2, VP3 and VP4 are colored in blue, green, red, and magenta, respectively. The density of inhibitor 11526092 is shown as a grey mesh. (B) A zoomed in view of the inhibitor 11526092 (grey) in the VP1 pocket (blue). (C) An enlarged view of inhibitor 11526092 fitted into the density (contour level 0.8) and the interfacing residues. (D) Comparison of the VP1s between EV-D68-inhibitor 11526092 complex (blue) and control (light purple). Large movements are highlighted by magenta dashes.

The methyl on the benzene ring of 11526092 is involved with multiple hydrophobic interactions with TRP-93, ILE-95, ILE-119, VAL-239, while the other analogs 11526091 and 11526093 have a nitro- or fluorine group at this position, respectively, which would either interfere or weaken this interaction. Among the three EV-D68 and inhibitor reconstructions, only the EV-D68-11526092 complexes shows clear densities for the inhibitor in the VP1 hydrophobic pocket region at a contour level of 0.8 (Fig. 3A-C), demonstrating good occupancy of 11526092. Based on the structure of EV-D68 (Fermon strain) complexed with pleconaril (PDB: 4wm7), 11526092 binds to the VP1 hydrophobic pocket in a similar manner with highly conserved interfacing residues (Fig. S9). However, comparison between the structures of EV-D68 MO in complex with 11526092 and the native EV-D68 MO strain shows conformational changes in the VP1 GH and EF loops (Fig. 3D). Residues 216-218 on the GH loop move more than 2 Å away from the hydrophobic pocket (based on the *Cα atoms*) (Fig.3D, Fig. S10). Similarly, obvious outward movements are observed on EF loop, residues 150-152 (Fig.3D, Fig. S10). Because the VP1 GH and EF loops are closely associated with the canyon region, these residue movements likely contribute to binding dynamics, affecting the area where receptor binding would occur and interfere with binding and cell tropism (*40*).

## Discussion

Recently, antiviral efforts towards the development of anti-picornaviral compounds have focused on the capsid-binding mechanism of action (*41*). The most advanced compound, pleconaril was tested on 215 clinical isolates of EV serotypes and demonstrated prommising activity (e.g. IC_50_ ≤ 0.03 μM for all isolates tested and IC_90_ inhibition of ~90% of isolates were ≤ 0.18 μM) (*21*). Pleconaril had a similar inhibitory effect on EV-71 both *in vitro* (IC_50_ 0.34-1.42 μM) and in the mouse model (*42*). However, results from different clinical trials of pleconaril were ambiguous despite the wide antiviral activity *in vitro*. Pleconaril proved useful in 2 of 3 neonates with severe enteroviral hepatitis in one study (*43*). It also resulted in clinical improvement in 78% of the patients with chronic meningoencephalitis (*43, 44*). A double-blind placebo-controlled trial on infants with EV meningitis also found no significant effect with pleconaril (*45*). It had no statistically significant effect on the treatment of the common cold at a dose of 400 mg given three times a day over five days, while adverse effects (headache and menstrual dysfunction) were common, causing some women to bleed between menstrual periods, interfering with hormonal birth control and leading to unintended pregnancy. Subsequent studies suggested that pleconaril induced CYP3A4, as this enzyme is known to be primarily responsible for the metabolism of birth control medication (*46, 47*). Viruses with resistance to pleconaril were also found in 10.7% of pleconaril-treated patients (*44, 48*). With the clinical failure of this and other related compounds, it is therefore imperative that we restart the antiviral pipeline to develop a treatment for the numerous diseases caused by EVs, RVs and CVs, as there is a high unmet need.

We have now described the development of a novel isoxazole-3-carboxamide pleconaril analog 11526092, which is a more potent inhibitor of CVB3 (Table 1) and EV-D68 (EC_50_ 59 nM for 11626092 versus 430 nM for pleconaril (*32*)). A cryo-EM structure shows that 11526092 is bound to the VP1 hydrophobic pocket of EV-D68 (MO strain) in a similar manner as pleconaril was previously shown in EV-D68 (Fermon strain) (*32*) with residues associated with the binding pocket being highly conserved (Fig. S9). In the previously reported structure of EV-D68 (Fermon) in complex with pleconaril, ILE211 on the VP1 GH loop shifts towards the pocket and thus likely locks the pocket (*32*). In contrast, 11526092 in EV-D68 (MO strain) pushes the VP1 GH loop and EF loop away from the pocket (Fig. 1D, Fig. S10). This was also observed for pleconaril in EV-D68 (MO strain) (Fig. S11, S12) suggesting both destabilize the protein and opening it up (*40*). Thus, it appears that pleconaril demonstrates different mechanisms towards EV-D68, stabilizing Fermon and destabilizing MO strains. These new cryo-EM structures provide atomic level information on the inhibition mechanism that may be beneficial for future EV-D68 antiviral drug design and suggest we will need to consider how molecules bind to the VP1 in different strains.

We have demonstrated the efficacy of 11526092 for treatment of an EV-D68 respiratory infection in four-week-old AG129 mouse model. The EV-D68 respiratory model based on intranasal infection of 4-week-old AG129 mice is a non-lethal model that includes viremia with rapid tissue distribution of virus, including high virus titers in lung tissues and an elevation of proinflammatory cytokines in the lung. Treatment with all doses of 11526092 significantly reduced blood virus (viremia) on days 1 and 3 post-infection. No virus was detected in the blood of animals treated on day 5 post-infection indicating virus clearance from the blood by this time. In addition, a dose of 30 mg/kg/day reduced lung virus titers on day 5 post-infection compared to placebo-treated mice. Interestingly, a full elimination of detectable viremia without full tissue-specific clearance is not unprecedented, even while providing full survival protection for other viruses (*49*). This would suggest that full viremia suppression may be a strong indicator of disease progression for viral infections related to human disease.

Production of inflammatory cytokines such as IL-1β, IL-6, and TNF-α is also important in control of respiratory infections. Cellular immunity was evaluated by quantitation of cytokines and chemokines in lung lavage samples collected on days 1, 3, and 5 p.i. using a multiplex ELISA. All doses of 11526092 significantly reduced lung concentrations of IL-3, IL-5, GM-CSF, and RANTES on day 1 after treatment and challenge infection. Additional reductions in cytokine concentrations after treatment and infection included IFNγ and MCP-1 on day 1 for the 10 and 100 mg/kg treatment groups, respectively. An increase in IL-1α, IL-1β, IL-4, and IL-6 were observed on day 3 for the 100 mg/kg treatment groups. In addition, an increase in IL-10 and MIP-1α were observed on day 5 for the 10 and 30 mg/kg treatment groups, respectively (Fig S5). Evaluating the efficacy of 11526092 administered therapeutically (after virus infection) in this model would be valuable to determine the window for administration of 11526092 in future.

In contrast, neurological studies demonstrated a lack of the prophylactic and therapeutic efficacy of 11526092 (dosed up to 10 mg/kg/day IP) in terms of survival, weight loss, neurological scores, and virus titers in the blood when treated at either 2-hour pre-infection or 4 hours post-infection. The positive control guanidine (100 mg/kg/day IP) was more effective in this model, but this is not considered a clinically viable option. The poor efficacy of 11526092 in the mouse model could be due to the level of exposure as demonstrated in our earlier *in vitro* ADME studies (e.g. poor mouse metabolic stability (Table 3)) and pharmacokinetics study in mouse (*28*) and this may require substantially higher concentrations when dosed IP. In humans pharmacokinetics may also be improved as 11526092 was considered stable when assessed for metabolic stability in human microsomes (Table 3). It was assumed that the previously demonstrated failure of pleconaril in the clinic and breakthrough pregnancies were due to CYP3A4 induction. As CYP3A4 induction is likely a key issue (*46, 47*) we previously tested 11526092 in cryopreserved human hepatocytes from a single donor in duplicate using rifampicin as positive control (*50*). At 1 μM pleconaril resulted in 1.7x induction (13% of control) vs 11526092 1.2x (3% of control). These results suggest 11526092 is less likely to be a CYP3A induction risk versus pleconaril. We have now demonstrated that neither 11526092 nor pleconaril is an agonist of the pregnane X-receptor (PXR) which is involved in induction of CYP3A4 and instead inhibiting CYP3A4 in human liver microsomes as both pleconaril and 11526092 demonstrate autoactivation at higher concentrations (Fig. S13). Whether these are clinically significant *in vitro* observations may be considered in future. In the case of 11526092, due to its more potent antiviral activity *in vitro*, this effect on CYP3A4 may be less pronounced than for pleconaril which has a reduced antiviral activity and will require a higher dose. 11526092 also has activity against pleconaril resistant viruses (*28*) which could suggest resistance to this compound may be less of an issue than for pleconaril.

In conclusion, 11526092 is a promising antiviral compound for multiple EVs, binding to EV-D68 in a similar manner to pleconaril, and we describe for the first time how such compounds may use unique mechanisms of action for different strains of the same virus. 11526092 is well tolerated in mice and demonstrates promising antiviral efficacy in the respiratory model of AFM. Future studies in the mouse neurological model at much higher doses are justified to assess efficacy. Based on this study and in the absence of any suitable treatments for EV-D68, future clinical testing of 11526092 may be considered for treating this virus.

## Acknowledgments

Dr. Eun-Chung Park, Dr. Mindy Davis and colleagues are kindly acknowledged for their support of this project and assistance with the NIAID virus *in vivo* testing and screening capabilities. Dr. Patricia Vignaux and Dr. Ana Puhl are acknowledged for careful reading. Dr. Michaela Schmidtke is acknowledged for our earlier work on this series of compounds.

## Funding

National Institutes of Health (National Institute of Neurological Disorders and Stroke (NINDS) grant 1R01NS102164-01 (SE)

National Institutes of General Medical Sciences grant: R44GM122196-02A1 (SE)

National Institute of Allergy and Infectious Diseases (NIAID) program for non-clinical and pre-clinical services (SE)

This work was supported R01-AI011219 (R.J.K), and by NIH/NIAID contract HHSN272201700060C (R.J.K; PI: K. Satchell).

Funding was provided by the National Institute of Health contract number HHSN27220100041I, Task Order A16, from the Virology Branch, Division of Microbiology and Infectious Diseases, National Institute of Allergy and Infectious Diseases, National Institute of Health, USA.

## Author contributions

Conceptualization: SE, VM

Methodology: JF, BS, BT, BLH, OR, EK, AE, PC, SL, JF, MR, KT, RJK

Investigation: TL, JF, BS, BT, BLH, OR, EK, AE, PC, SL, JF, MR, KT, TK, RJK, VM, SE

Visualization: TL, JF

Funding acquisition: SE, VM

Project administration: SE

Supervision: SE, VM, RJK

Writing – original draft: SE, VM

Writing – review & editing: SE, TRL, VM

## Data and materials availability

“All data are available in the main text or the supplementary materials. Cryo-EM files have been submitted to the PDB.”

