## Supplemental materials for "Strain dependent structural effects and *in vivo* efficacy of enterovirus-D68 inhibitors"

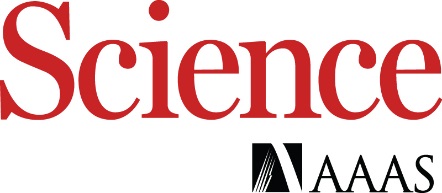

Supplementary Materials for

**Strain dependent structural effects and *in vivo* efficacy of enterovirus-D68 inhibitors**

Authors: Thomas Lane^1*^, Jianing Fu^2*^, Barbara Sherry^3^, Bart Tarbet^4,5^, Brett L. Hurst^4,5^, Olga Riabova^6^, Elena Kazakova^6^, Anna Egorova^6^, Penny Clarke^7^, J. Smith Leser^7^, Joshua Frost^8^, Michael Rudy^7^, Ken Tyler^7-9^, Thomas Klose^2^, Richard J. Kuhn^2^, Vadim Makarov^6^ and Sean Ekins^1†^

**This PDF file includes:**

Supplementary Text

Supplementary Materials and Methods

Supplementary References

Figs. S1 to S17

Tables S1 to S6

**Supplementary Text**

Production of inflammatory cytokines such as IL-1β, IL-6, and TNF-α is also important in control of respiratory infections. Cellular immunity was evaluated by quantitation of cytokines and chemokines in lung lavage samples collected on days 1, 3, and 5 p.i. using a multiplex ELISA. Fig. S4 shows concentrations of IL-1α, IL-1β, IL-2 and IL-3 in lung homogenates from mice following treatment with 11526092 and EV-D68 infection. Treatment with 11526092 at a dose of 30 mg/kg/day increased concentrations of IL-1α and IL-1β on day 3 p.i. compared to placebo-treated mice. In addition, the 30 mg treatment also increased IL- 3 on day 3 p.i. (Fig. S4). In addition, all doses of 11526092 decreased IL-3 on day 1. Concentrations of IL-4, IL-5, IL-6, and IL-10 in lungs from mice following treatment with 11526092 and EV-D68 infection were also evaluated. Treatment with 11526092 at a dose of 100 mg/kg/day increased concentrations of IL-4, IL-5, and IL-6 (Fig. S4) on day 3 p.i. compared to placebo-treated mice. However, the 100 mg dose also decreased IL-5 on day 1 p.i.. In addition, all doses of 11526092 decreased IL-5 on day 1 p.i., but the 30 mg treatment increased IL-6 on day 3 p.i.. Concentrations of IL-12p70, IL-17, MCP-1 and IFNγ in lungs from mice following treatment and infection were additionally tested. Treatment with 11526092 at a dose of 100 mg/kg/day decreased concentrations of MCP-1 on day 1 p.i. and the 10 mg dose decreased IFNγ on day 1 p.i.. Other changes included guanidine treatment increasing IL-12p70 on day 3 p.i. and decreasing MCP-1 on days 1 and 3 p.i. Finally, concentrations of TNFα, MIP-1α, GM-CSF, and RANTES were assessed in lungs from mice following treatment and infection. Treatment with 11526092 at a dose of 30 mg/kg/day increased concentrations of MIP-1α on day 5 p.i. and all doses of 11526092 decreased GM-CSF and RANTES on day 1 p.i.. Other observations included guanidine treatment increasing MIP-1α on day 5 p.i. and decreasing GM-CSF and RANTES on day 1 p.i.

Cryo-EM:

For the reconstructions of EV-D68 MO in complex with 11526091 and 11526093, there is only weak density in the VP1 hydrophobic pocket region, indicating a low occupancy or no binding in (Fig. S14). This is more pronounced for 11526093 than for 11526091, suggesting that the former binds slightly stronger than the latter. In addition, compared to the control, no structural changes are observed in EV-D68 complexes with the two compounds (Fig. S15). However, a drop in particle densities still indicates a possibility that both molecules inhibit the virus using different mechanisms. Moreover, the reconstruction of EV-D68 MO in complex with pleconaril shows a similar conformational change as EV-D68 MO in complex with inhibitor 11526092, which is different to the movement observed in EV-D68 Fermon in complex with pleconaril (Fig. S10-S12), indicating that the same compound may have different anti-viral mechanisms among viral strains. An example of typical map densities (VP2 and VP3 sections) at 2 Å resolution are shown in Fig. S16 for reference. Estimates of map resolution are also shown in Fig. S17.

Supplementary Materials and Methods

Ethics statement

Ethics regulation of laboratory animals: This study was conducted in accordance with the approval of the Institutional Animal Care and Use Committee of Utah State University dated March 2, 2019 (expires March 1, 2022). The work was done in the AAALAC-accredited Laboratory Animal Research Center of Utah State University. The U. S. Government (National Institutes of Health) approval was renewed March 2, 2019 (PHS Assurance No. D16-00468[A3801-01], expires March 1, 2022) in accordance with the National Institutes of Health Guide for the Care and Use of Laboratory Animals (Revision; 2011).

Chemicals and reagents

All reagents and solvents were purchased from commercial suppliers and used without further purification. ^1^H Spectra were measured on Bruker AC-300 (300 MHz, ^1^H). Chemical shifts were measured in DMSO-d_6_ or CDCl_3_, using tetramethylsilane as an internal standard, and reported as units (ppm) values. The following abbreviations are used to indicate the multiplicity: s, singlet; d, doublet; t, triplet; m, multiplet; dd, doublet of doublets; brs, broad singlet; brm, broad multiplet. Mass spectra were recorded on Finnigan MAT INCO 50 mass spectrometer (EI, 70 eV) with direct injection. The purities of the final compounds were analyzed on an Agilent 1290 Infinity II HPLC system coupled to Agilent 6460 triple-quadrupole mass spectrometer equipped with an electrospray ionization source. The chromatographic separation was carried out on an Agilent Eclipse Plus C18 RRHD column (2.1 × 50 mm, 1.8 µm) at 40 ºC; sample injection volume was 0.2 µL. The mobile phase comprising 0.1 % formic acid/water (A), and 0.1 % formic acid and 85 % acetonitrile / water (B) was programmed with gradient elution (0.0-3.0 min, 60 % B; 3.0-4.0 min, 60 % to 97 % B; 4.0-6.0 min, 97 % B; 6.0-6.1 min, 97 % to 60 % B) at a flow rate of 0.4 mL/min. The mass spectrometric detection was operated in positive ion mode. Optimal parameters were: capillary voltages of 3500 V, a nebulizer pressure of 35 psi, a gas temperature of 350 ºC, a gas flow rate of 12 L/min.

Pleconaril was synthesized according to a previous publication (*1*) and its analytical data was identical to this published information. All final compounds were > 99 % pure.

### Compounds synthesis

3-(3-Methyl-4-(3-(3-*N*,*N*-dimethylcarbamoyl-isoxazol-5-yl)propoxy)phenyl)-5-trifluoromethyl-1,2,4-oxadiazole **11526092** was synthesized in 5 steps, and complete experimental protocols have been described by Egorova *et al*. (*2*). According to Scheme 1, 4-cyanophenol is *O*-alkylated with chloropentyne in the presence of potassium carbonate and potassium iodide. The obtained pentynyloxy-benzonitrile is treated with an excess of hydroxylamine in alcohol to provide the corresponding hydroxamic acid derivative. Its acylation with trifluoroacetic anhydride in pyridine leads to the cyclization of 1,2,4-oxadiazole and the formation of ((pentynyloxy)phenyl)-5-trifluoromethyl-1,2,4-oxadiazole. This key intermediate reacts with ethyl 2-chloro(hydroxyimino)acetate to form the 3-carbethoxy-isoxazole derivative. Finally, the substitution of the 3-carbethoxy group using a dimethylamine solution in dioxane gives the appropriate dimethylcarbamoyl derivative **11526092**. White solid, yield 70 %, mp 88-89 ºС (EtOH/H_2_O). MS (EI; rel. int., %): m/z 424.3 (M^+^, 37). ^1^H NMR (DMSO-d_6_; δ, ppm): 2.22 (2H, dt, *J* = 6.5, 7.6, CH_2_CH_2_CH_2_O), 2.24 (3H, s, CH_3_Ph), 3.00 (3H, s, NCH_3_), 3.05 (2H, t, *J* = 7.6, CH_2_CH_2_CH_2_O), 3.10 (3H, s, NCH_3_), 4.20 (2H, t, *J* = 6.5, CH_2_CH_2_CH_2_O), 6.50 (1H, s, isoxazole), 7.20 (1H, d, *J* = 8.8, H6), 7.80 (1H, d, *J* = 1.5, H5), 7.81 (1H, dd, *J* = 1.5, 8.8, H3). Anal. calcd for C_19_H_19_F_3_N_4_O_4_: C, 53.77; H, 4.51; N, 13.20. Found: С, 53.78; Н, 4.50; N, 13.22.The yields of all chemical steps are high, and the final compound can be readily synthesized in large quantities.

Other analogs **11526091** and **11526093** were synthesized in a similar manner.

3-(3-Nitro-4-(3-(3-*N*,*N*-dimethylcarbamoyl-isoxazol-5-yl)propoxy)phenyl)-5-trifluoromethyl-1,2,4-oxadiazole **11526091**. White solid, yield 37 %, mp 62-64 ºС (EtOH). MS (EI; rel. int., %): m/z 455.3 (M^+^, 52). ^1^H NMR (DMSO-d_6_; δ, ppm): 2.20 (2H, dt, *J* = 6.5, 7.6, CH_2_CH_2_CH_2_O), 2.98 (3H, s, NCH_3_), 3.00 (2H, t, *J* = 7.6, CH_2_CH_2_CH_2_O), 3.05 (3H, s, NCH_3_), 4.40 (2H, t, *J* = 6.5, CH_2_CH_2_CH_2_O), 6.51 (1H, s, isoxazole), 7.55 (1H, d, *J* = 8.8, H6), 8.32 (1H, dd, *J* = 1.5, 8.8, H5), 8.50 (1H, d, *J* = 1.5, H3). Anal. calcd for C_18_H_16_F_3_N_5_O_6_: C, 47.48; H, 3.54; N, 15.38. Found: С, 47.49; Н, 3.53; N, 15.40.

3-(3-Fluoro-4-(3-(3-N,N-dimethylcarbamoyl-isoxazol-5-yl)propoxy)phenyl)-5-trifluoromethyl-1,2,4-oxadiazole **11526093**. White solid, yield 42 %, mp 83-85 ºС (hexane). MS (EI; rel. int., %): m/z 428.3 (M^+^, 29). ^1^H NMR (DMSO-d_6_; δ, ppm): 2.21 (2H, dt, *J* = 6.5, 7.6, CH_2_CH_2_CH_2_O), 2.99 (3H, s, NCH_3_), 3.10 (3H, s, NCH_3_), 3.05 (2H, t, *J* = 7.6, CH_2_CH_2_CH_2_O), 4.25 (2H, t, *J* = 6.5, CH_2_CH_2_CH_2_O), 6.52 (1H, s, isoxazole), 7.40 (1H, t, *J* = 8.3, H3), 7.80-7.90 (2H, m, H5, H6). Anal. calcd for C_18_H_16_F_3_N_4_O_4_: C, 50.47; H, 3.77; N, 13.08. Found: С, 50.49; Н, 3.73; N, 13.00.

**Scheme 1. Five-step synthesis of 11526092**

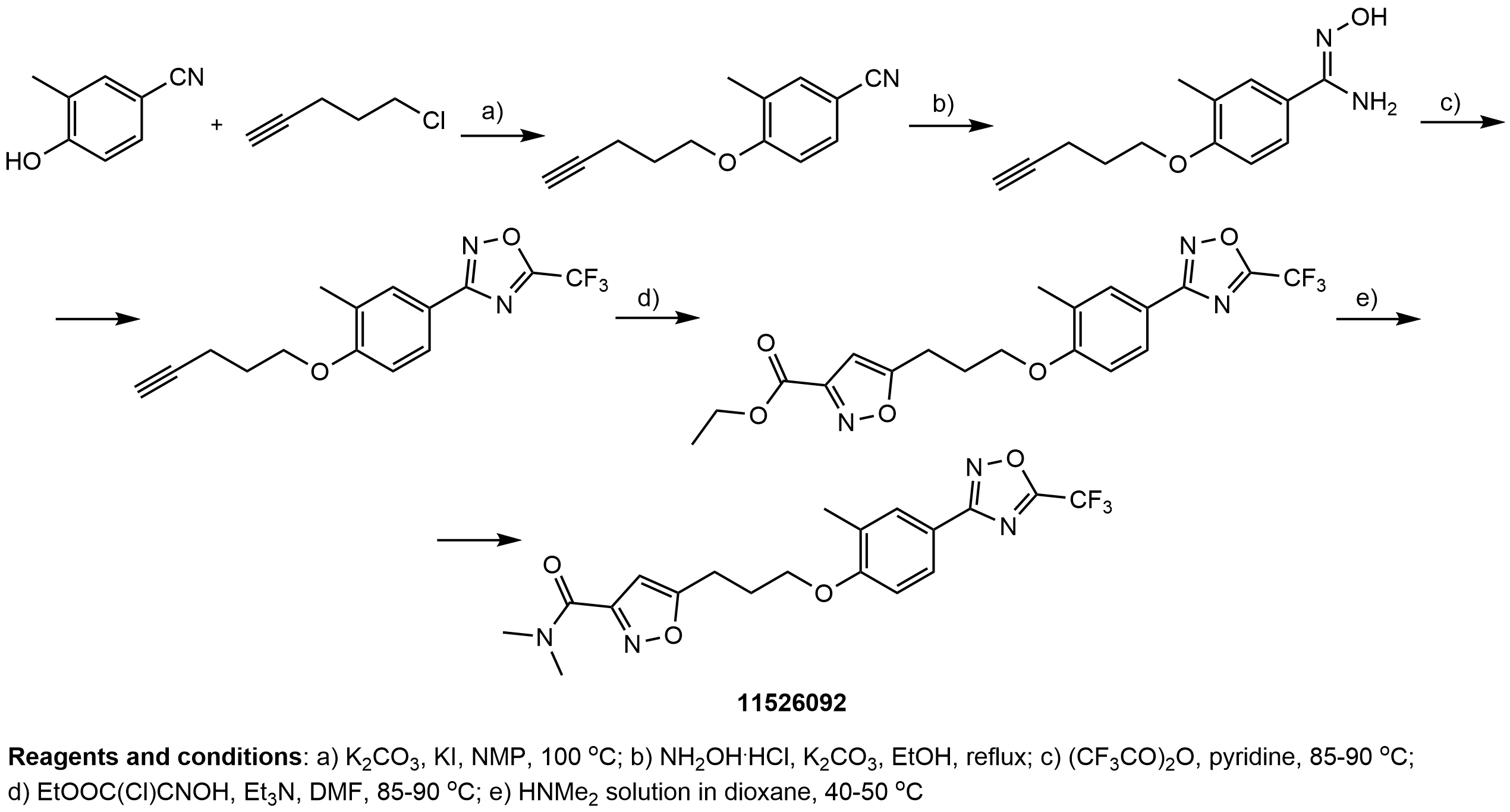

*In vitro* antiviral assays

*CVB3:* Drugs were solubilized in DMSO and then serially diluted 2-fold in cell culture media and then drugs or DMSO vehicle were mixed with sufficient CVB3 for an MOI of 1 or 3 PFU per cell and used to immediately infect HeLa cell cultures in 96-well plates. After incubation for 48 hours, cell monolayers were stained with crystal violet, the optical density determined relative to DMSO-treated controls, and the Inhibitory Concentration 50% (IC_50_) calculated. N = 6 wells per case (e.g. dilution). In addition, the Cytotoxicity Concentration 50% (CC_50_) was assessed for each virus (using the same protocol in the absence of virus) and determined to be >100 µM (not tested at higher concentrations).

For low MOI infections, HeLa cell cultures in 96-well plates were infected with CVB3 at a low MOI such that only a fraction of cells was infected and then incubated for 6 hours to allow ample time for virus entry and establishment of infection. Two-fold serial dilutions of drug (as above) were added to the culture wells. After incubation for 48 hours to allow virus spread, cell monolayers were stained with crystal violet, the optical density determined relative to DMSO-treated controls, and the Inhibitory Concentration50 (IC_50_) calculated. N = 6 wells per case.

EVD68: Human rhabdomyosarcoma (RD) cells were treated with Pleconaril or indicated analogs (25 µM) for 24 hours prior to infection with EVD68, strain IL/14-18952 (IL52), at a multiplicity of infection of 0.1. At 24, 48, and 72 hours following infection cells were harvested and total RNA was prepared using a Qiagen RNeasy Plus Micro Kit with RNA carrier (Hilden, GER) and used to make cDNA using a Bio-Rad iScript RT supermix (Berkley, CA). Equivalent volumes of cDNA were used for PCR with degenerate primers targeting the VP1 gene (*3*). Samples were compared to a plasmid standard curve.

Vero 76 cells Reduction of virus-induced cytopathic effect (Primary CPE assay)

Confluent or near-confluent cell culture monolayers of Vero 76 cells are prepared in 96-well disposable microplates the day before testing. Cells are maintained in MEM supplemented with 5% FBS. For antiviral assays the same medium is used but with FBS reduced to 2% and supplemented with 50 µg/ml gentamicin. Compounds are dissolved in DMSO, saline or the diluent requested by the submitter. Less soluble compounds are vortexed, heated, and sonicated, and if they still do not go into solution are tested as colloidal suspensions. The test compound is prepared at four serial log_10_ concentrations, usually 0.1, 1.0, 10, and 100 µg/ml or µM (per sponsor preference). Lower concentrations are used when insufficient compound is supplied. Five microwells are used per dilution: three for infected cultures and two for uninfected toxicity cultures. Controls for the experiment consist of six microwells that are infected and not treated (virus controls) and six that are untreated and uninfected (cell controls) on every plate. A known active drug is tested in parallel as a positive control drug using the same method as is applied for test compounds. The positive control is tested with every test run.

Growth media is removed from the cells and the test compound is applied in 0.1 ml volume to wells at 2X concentration. Virus, normally at ~60 CCID_50_ (50% cell culture infectious dose) in 0.1 ml volume is added to the wells designated for virus infection. Medium devoid of virus is placed in toxicity control wells and cell control wells. Plates are incubated at 37 ^o^C with 5% CO_2_ until marked CPE (>80% CPE for most virus strains) is observed in virus control wells. The plates are then stained with 0.011% neutral red for approximately two hours at 37^o^C in a 5% CO_2_ incubator. The neutral red medium is removed by complete aspiration, and the cells may be rinsed 1X with phosphate buffered solution (PBS) to remove residual dye. The PBS is completely removed, and the incorporated neutral red is eluted with 50% Sorensen’s citrate buffer/50% ethanol for at least 30 minutes. Neutral red dye penetrates living cells, thus, the more intense the red color, the larger the number of viable cells present in the wells. The dye content in each well is quantified using a spectrophotometer at 540 nm wavelength. The dye content in each set of wells is converted to a percentage of dye present in untreated control wells using a Microsoft Excel computer-based spreadsheet and normalized based on the virus control. The 50% effective (EC_50_, virus-inhibitory) concentrations and 50% cytotoxic (CC_50_, cell-inhibitory) concentrations are then calculated by regression analysis. The quotient of CC_50_ divided by EC_50_ gives the selectivity index (SI) value. Compounds showing SI values ≥10 are considered active.

Vero 76 cells Reduction of virus yield (Secondary VYR assay)

Active compounds are further tested in a confirmatory assay. This assay is set up like the methodology described above only eight half-log_10_ concentrations of inhibitor are tested for antiviral activity and cytotoxicity. After sufficient virus replication occurs (3 days for SARS-CoV-2), a sample of supernatant is taken from each infected well (three replicate wells are pooled) and tested immediately or held frozen at -80 °C for later virus titer determination. After maximum CPE is observed, the viable plates are stained with neutral red dye. The incorporated dye content is quantified as described above to generate the EC_50_ and CC_50_ values. The VYR test is a direct determination of how much the test compound inhibits virus replication. Virus yielded in the presence of test compound is titrated and compared to virus titers from the untreated virus controls. Samples were collected 3 days after infection. Titration of the viral samples (collected as described in the paragraph above) is performed by endpoint dilution (*4*). Serial 1/10 dilutions of virus are made and plated into 4 replicate wells containing fresh cell monolayers of Vero 76 cells. Plates are then incubated, and cells are scored for presence or absence of virus after distinct CPE is observed (3 days after infection), and the CCID_50_ calculated using the Reed-Muench method (*4*). The 90% (one log_10_) effective concentration (EC_90_) is calculated by regression analysis by plotting the log_10_ of the inhibitor concentration versus log_10_ of virus produced at each concentration. Dividing EC_90_ by the CC_50_ gives the SI value for this test.

*In Vitro* ADME assays

*In vitro* ADME studies described below were performed by BioDuro (San Diego, CA).

Bioanalytical Method for *In Vitro* ADME Studies

Test compounds were analyzed by reverse phase HPLC with a Kinetex 2.6µ C18 100A column (3.0 mm x 50 mm, Phenomenex (Torrance Ca)) using Shimadzu (Columbia, MD) LC-20AD system. The mobile phase consisted of solvent A (Water with 0.1% Formic) and solvent B (acetonitrile with 0.1% Formic). The MS detection was performed by using an API 4000 Q trap system. The amount of parent compound was determined on the basis of the peak area ratio (compound area to internal standard area).

Kinetic Solubility

396 µL of Universal Aqueous buffer (pH 7.4), FaSSIF-V2 (pH 6.5) or FaSSGF (pH 1.6) was added to 4 µL of a 50 mM DMSO stock solution of 11526902. Wells were agitated for 4 h at room temperature (RT), then let stand for 30 min at RT followed by filtration. The compound was then diluted to serial concentrations with DMSO, followed by serial dilutions with ACN: H_2_O (1:1) prior to LCMS analysis. The calculated concentration (µM) of soluble 11526092 was determined in reference to a standard curve.

Caco-2 Permeability

Caco-2 cells (ATCC, Manassas, VA) were grown on 24-well (pore size: 0.4 µm) polycarbonate filters. The monolayers were pre-incubated with pre-warmed HBSS (Hank’s balanced salt solution) containing 2.5% HEPES buffer (pH 7.4) for 0.5 h at 37ºC. After pre-incubation, the buffer was removed and 11526092 was added to reach a final concentration of 10 µM. 2% bovine serum albumin (BSA) was added to the receiver buffer for the study. The total volume was 400 µL for the apical (A) side and 1200 µL for the basolateral (B) side. For apical to basolateral transport study (A-B), 100 µL each was collected from both sides for sample analysis at the start of the assay and then 200 µL was collected from the apical side at 90 minutes (end of the study). The same timepoints and amounts were used for the basolateral to apical transport study (B-A).

The apparent permeability coefficient (Papp) was calculated from the following equation.

Where:

v = Volume of the receiver cell

A = Exposed surface area (0.64 cm^2^)

C = Initial donor concentration

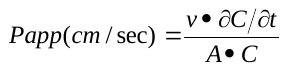

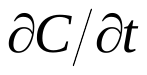
= Change in receiver concentration over time

Human CYP Inhibition

Human liver microsome solution (0.2 mg/ml) (Sekisui Xenotech, Kansas City, KS), along with substrate, was aliquoted into a 0.05 M phosphate buffer (pH=7.4) in 1.1 ml tubes. Study samples (containing either control inhibitor or test compound) were into added into the tubes, vortexed gently and pre-incubated for 5 min at 37°C. 20 µL of NADPH solution was aliquoted into all tubes, then vortexed to start the reaction and to assure adequate mixture of the NADPH. After mixing, the tubes were incubated for 20 min at 37°C in a shaking water bath and then quenched in 300 ml formic acid/acetonitrile solution. After quenching the samples were vortexed vigorously for 1 min and centrifuged at 4,000 rpm for 15 min (4°C). 100 µL of supernatant was transferred to 0.65 ml tubes, for LCMS analysis by the bioanalytical method described earlier. The CYP450 substrates and control inhibitors for each enzyme was as follows: CYP1A2 (phenacetin, naphthoflavone), CYP2C9 (diclofenac, sulfaphenazole), CYP2C19 (omeprazole, tranylcypromine), CYP2D6 (dextromethorphan, quinidine), CYP3A4 (midazolam, ketoconazole).

Liver microsome stability

Liver microsome solutions (197.5 µL, 1 mg/ml protein concentration) (Human (BioIVT; Catalog: X008070) and mouse (BioIVT; Catalog:; X008070) microsomes (Westbury, NY)) were aliquoted into 1.1 ml tubes, to which 2.5 µL of positive control and 11526092 stock solutions (100 µM in DMSO) were added. The tubes were vortexed gently, pre-incubated for 5 min at 37°C, then 50 µL of 5 mM NADPH or LM buffer (no NADPH buffer) was added into the tubes. For analysis, an aliquot of 15 µL was removed from each tube at 0, 5, 15, 30 and 60 min (without-NADPH reaction:0, 30 and 60 min) and quenched with 300 µL of 25 ng/ml propranolol in acetonitrile. Samples were vigorously vortexed for 1 min and then centrifuged at 4,000 rpm for 15 min at 4 ̊C. 100 µL of supernatant from each sample was transferred to 0.65 ml tubes for LCMS analysis. The amount of parent compound was determined based on the peak area ratio (compound area to IS area) for each time point. Clearance rates were calculated by the equation:

CL_int_ (µL/min/mg protein) = Ln (2)*1000 /T_1/2_ / Protein Conc.

Protein Binding in Plasma

The donor side of dialysis inserts were filled with 200 μL plasma (human (Cat#HUMANPLK21804231) and mouse (MSE00PLK2YNN) plasma from BioIVT, (Westbury, NY)) containing 5 μM drug and 0.5% of DMSO and the receiver side of the dialysis inserts was filled with 350 μL of PBS buffer (100 mM, pH 7.4). The prepared dialysis apparatus was placed in a shaker (37 ̊C, 100 rpm) for 5 hours. Two tubes with plasma containing 5 μM 11526092 were also prepared for stability test, one tube was placed in the freezer (4 ̊C) for 5 h, and the other tube was placed in shaker (3 ̊C, 100 rpm) for 5 h. Samples were collected from the donor and receiver sides of each dialysis insert. The same volume of blank plasma was added to buffer samples and blank buffer to plasma samples to make sure all sample mixtures contain 50% plasma and 50% buffer. 50 µL of each sample was mixed with 300 μL of acetonitrile containing 25 ng/ml internal standard (propranolol). All samples were vortexed for 1 minute and then centrifuged at 4000 rpm, 4℃ for 15 min. 100 µL of the supernatant was transferred to 0.65 ml tube for LCMS analysis. The amount of compound was determined on the basis of the peak area ratio (compound area to internal standard area) for the two sides, and protein binding is determined using the following equations: %Bound = 100 x ([Area Ratio of Donor] 5h *5 - [Area Ratio of Receiver] 5h) / ([Area Ratio of Donor] 5h *5). The percentage remaining at 37 ℃ after 5h was calculated on the basis of the amount measured at 0 ℃ after 5h.

PXR and hERG inhibition

PXR agonist activity and hERG inhibition were tested by the ThermoFisher Scientific SelectScreen^TM^ Profiling Service (Life Technologies Corporation, (Chicago, IL 60693). The PXR assay uses a LanthaScreen TR-FRET competitive binding assay and used SR-12813 as a positive control. a Tb-labeled anti-GST antibody is used to indirectly label the receptor by binding to its GST tag. Competitive ligand binding is detected by a test compound’s ability to displace a fluorescent ligand (tracer) from the receptor, which results in a loss of FRET signal between the Tb-anti-GST antibody and the tracer. The hERG screening used a fluorescence polarization assay previously described (*5*) and E-4031 as a positive control.

Maximum tolerated dose studies

Maximum tolerated dose studies were performed for 10-day and 4-week-old AG129 mice. For each age group a total of 20 mice were randomized into four groups of 5 mice each enabling 3 different doses for each (Table S2). Mice were treated twice daily for five days with 11526092 in a suspension of 0.5% carboxymethylcellulose (CMC) and 0.5% Tween 20 in sterile water per os (p.o.). Mice were weighed prior to treatment and then daily thereafter to assess the effects of treatment on body weight. All mice were observed for morbidity and mortality through day 14.

*In vivo* efficacy assay: Respiratory model

Using previously described methods (*6*) four-week-old male and female AG129 mice from a specific-pathogen-free colony maintained at the Utah Science Technology and Research (USTAR) building at Utah State University (USU) for this experiment. The mice were bred and maintained on irradiated Teklad Rodent Diet (Harlan Teklad) and autoclaved tap water at the USTAR building of USU. 11526092 was prepared in 0.5% carboxymethylcellulose (CMC) and 0.5% Tween 20 in sterile water. Guanidine HCl (guanidine) was obtained from Sigma-Aldrich (St. Louis, MO) and served as a positive control at a dose of 100 mg/kg/day. Sterile saline was used as a vehicle for guanidine. EV D68 was obtained from BEI Resources, NIAID, NIH: Enterovirus D68, US/MO/14-18949, NR-49130. The virus was serially passaged 30 times in the lungs of 4-week-old AG129 mice and then plaque-purified three times in Rhabdomyosarcoma (RD) cells obtained from the American Type Culture Collection (Manassas, VA). The resulting virus stock was amplified twice in RD cells to create a working stock. The virus used for infection was designated EV-D68 mouse passage 30 (MP30), plaque purified (PP). A total of 65 mice were randomized into 5 groups of 12 mice each and one group of 6 mice (normal controls) (Table S3). Mice were treated per os (PO) with 11526092 once daily for 5 days beginning 2 hours pre-infection. Treatment with guanidine started 4 hours post-infection and continued twice daily for 5 days. The placebo (vehicle only) was administered by p.o. once daily for 5 days starting 2 hours pre-virus exposure. Mice were anesthetized by i.p. injection of ketamine (100 mg/kg) and then infected via intranasal (IN) instillation of 1 x 104.5 CCID50 of EV-D68 MP30 PP in a 90 μl volume of MEM. Mice were weighed prior to treatment and then daily thereafter to assess the effects of treatment on ameliorating weight loss due to virus infection. Four mice from each treatment group were euthanized on days 1, 3, and 5 post-infection for evaluation of blood and lung virus titers, and lung cytokine concentrations. In addition, one mouse from the normal controls was euthanized on days 1, 3, and 5 post-infection as a negative control for cytokine analysis. Viremia was evaluated in whole blood on human rhabdomyosarcoma (RD) cells in microtiter plates, as described previously (*6, 7*). In addition, each mouse lung was homogenized in minimal essential media (MEM) solution and assayed for infectious virus in RD cells (*6, 7*). Fifty percent cell culture infectious doses (CCID50) were converted to CCID50 per gram of lung tissue prior to statistical analysis. Virus titer differences were evaluated by analysis of variance (ANOVA) on log-transformed values assuming equal variance and normal distribution. Following ANOVA individual treatment values were compared to placebo control by Dunnett’s pair-wise comparison test using Prism 9.1. Lung cytokine/chemokine determinations: samples (200 μl) from lung homogenates were tested for cytokines and chemokines using a chemiluminescent ELISA-based assay according to the manufacturer’s instructions (Quansys Biosciences Q-PlexÔ Array, Logan, UT). The Quansys multiplex ELISA is a quantitative test in which 16 distinct capture antibodies have been applied to each well of a 96-well plate in a defined array. Each sample supernatant was tested at 2 dilutions for the following: IL-1α, IL-1β, IL-2, IL-3, IL-4, IL-5, IL-6, IL-10, IL-12p70, IL-17, MCP-1, IFN-γ, TNFα, MIP-1α, GM-CSF, and RANTES. Cytokine and chemokine titers are reported in pg/ml of lung lavage fluid. Titer difference were evaluated by ANOVA on values assuming equal variance and normal distribution. In addition, treatment group mean values were evaluated by two-way ANOVA for effects based on the day post-infection using Prism 9.1. Mean body weights were analyzed by one-way analysis of variance (ANOVA) followed by Dunnett’s multiple comparison tests using Prism 9.1 (GraphPad Software Inc.). For each day post-infection, lung and blood virus titers from treated groups were compared to lung and blood titers from placebo-treated mice using a one-way analysis of variance (ANOVA) followed by Tukey’s pair-wise comparison test. For each cytokine/chemokine, the concentrations from treated mice were compared to placebo treated mice using a two-way ANOVA for effects based on the day post-infection.

*In vivo* efficacy assay: neurological model

Animals: 10-day-old AG129 mice from a specific-pathogen-free colony maintained at the Utah Science Technology and Research (USTAR) building at Utah State University. The mice were bred and maintained on irradiated Teklad Rodent Diet (Harlan Teklad) and autoclaved tap water at the USTAR building of Utah State University. 11526092 was prepared in 0.5% carboxymethylcellulose (CMC) and 0.5% Tween 20 in sterile water. Using previously described methods (*7*) two antiviral efficacy studies evaluating 11526092 were completed (Table S4 and S5). One study had antiviral drug treatment begin 4 hours post-infection, and the second study had the drug treatment begin 2 hours pre-infection. A total of 42 mice were randomized into five groups of 8 mice each as shown in Tables 1 and 2. Mice were treated twice daily for five days with 11526092 in a suspension of 0.5% carboxymethylcellulose (CMC) and 0.5% Tween 20 in sterile water per os (p.o.). Mice were weighed prior to treatment and then daily thereafter to assess the effects of treatment on body weight. In addition, all mice were observed for morbidity, neurological scores and mortality through day 14. Statistical analysis: Kaplan-Meier survival curves were generated and compared by the Log-rank (Mantel-Cox) test followed by pairwise comparison using the Gehan-Breslow-Wilcoxon test in Prism 8.4 (GraphPad Software Inc., La Jolla, CA). The mean body weights were analyzed by analysis of variance (ANOVA) followed by Tukey’s multiple comparison test using Prism 8.4. Neurological scores were graphed and evaluated using a Kruskal-Wallis test with a Dunn’s post-test for statistical significance compared to placebo-treated mice. Tissue virus titers were compared by a one-way ANOVA for each day with treated groups compared to placebo-treated mice.

Cryo-EM

Cells and viruses: Human Rhabdomyosarcoma cells (RD cells, CCL-136, American Type Culture Collection) were grown in medium composed of Dulbecco’s Modified Eagle Medium (DMEM, Sigma-Aldrich), 10% heat inactivated fetal bovine serum (HI-FBS, Sigma-Aldrich) together with 1X of nonessential amino acid (NEAA, Life Technologies). EV-D68 US/MO/14-18947 (EV-D68 MO) (GenBank accession no. AIS73051.1) was obtained from BEI Resources, National Institute of Allergy and Infectious Diseases, National Institute of Health. After passage in RD cells the virus was frozen and stored at -80℃.

VVirus propagation and purification: The EV-D68 MO used in structural studies was prepared using the following methods. RD cells were infected with EV-D68 MO with a multiplicity of infection of around 0.01. Two days post infection, the cells and supernatant were collected and spun down using a JA-10 rotor at 9,500 RPM. The cell pellets were frozen and thawed multiple times and spun down to remove cell debris. The combined supernatants were pelleted using a Ti 50.2 rotor at 48,000 RPM. The pellets were then incubated and resuspended in 250 mM HEPES, 250 mM NaCl buffer (pH=7.5) and 5mM (final concentration in the suspension) MgCl2 solution, 0.01mg/ml DNAse (Sigma-Aldrich), 0.8 mg/ml trypsin, 15mM EDTA, 1% (w/v) n-lauryl-sarcosine solution was added. Then the sample was pelleted down using a Ti 50.2 rotor at 48,000 RPM, resuspended and applied onto a potassium tartrate gradient (10%-40%, w/v) for the last round of ultracentrifugation using an SW41 rotor at 36,000 RPM. The purified virus sample was observed as a dominant blue band at the middle to lower part of the tube and was extracted and buffered exchanged to remove potassium tartrate.

Cryo-Sample preparation: For each of the inhibitors (11526091, 11526092 and 11526093) a 10mg/ml stock solution was made by dissolving the powder in DMSO. The three stock solutions were further diluted to 0.5mg/ml in PBS. Purified EV-D68 particles (approximately 1mg/ml in 20mM Tris, 120mM NaCl, pH=8.0 buffer) were mixed with each inhibitor at a molar ratio of 1:1500. After incubating for 6 hours at 4 ℃, for each of the virus-inhibitor mixtures and a control (EV-D68 MO alone), 3.5 µL solution was added to a glow-discharged 400 mesh lacey carbon film copper grid (Ted Pella Inc.). All samples were blotted for 4 seconds at 75%-80% humidity, and plunge frozen (Cryoplunge 3 system, Gatan) in liquid ethane.

 Cryo-EM and data collection: The Cryo-EM datasets were collected using two 300 kV Titan Krios Cryo-Transmission Electron Microscope (Thermo Fisher Scientific). Movies were collected using EPU or Leginon (*8*)with a K3 Direct Detection Camera (Gatan) at a magnification of 64,000X, resulting in a super resolution pixel size of 0.666 or 0.664 Å, with a defocus range from 0.4 to 2 µm (Table S6). A total electron dose of 36.06 or 36.3 electrons/Å^2^ over 2.6 seconds of exposure was recorded over 40 frames. Overall, 10,095 movies, 12,366 movies, 5,677 movies, 6789 movies and 3,972 movies were acquired for EV-D68 in complex with 11526091, 11526092, 11526093, pleconaril and control, respectively.

IImage processing, model building and refinement: For all datasets, motion correction, contrast transfer function (CTF), particle picking and extraction, 2-dimensional (2D) classification, 3D reconstruction and refinement were done using cryoSPARC (*9*). The final resolutions for all maps were estimated based on a gold-standard Fourier shell correlation cutoff of 0.143 (*10*). For model building and refinement, the same methods described below were applied for all datasets. The cryo-EM model of EV-D68 MO (PDB code: 6CSG) was selected as a template for model building and was manually fitted into the density maps using the program Chimera (*11*). Based on the initial fittings, the models were rebuilt in COOT (*12*)and processed in PHENIX (*13*) for real space refinements. The final validation of the atomic models was done through MolProbity (*14*), Chimera (*11*), COOT (*12*) and CCP4i2-PISA (*15*), which were used to visualize and determine the residues associated with the bound inhibitors. Refinement statistics are summarized in Table S6.

Docking

Compounds were docked in the various crystal structures that were previously published (PDB codes noted in legends) before our own cryo-EM work. The docking sphere was centralized from the position of the crystallized ligand with the default diameter (~10 Å depending on ligand shape). CHARMm based docking (CDOCKER) (*16*) was used to generate a maximum of 10 poses of each compound within this sphere using rigid docking. CDOCKER, a simulated annealing-based docking algorithm, was run within Discovery Studio (Biovia, San Diego, CA) using the following default parameters: 1000 dynamic steps, which included electrostatic interactions, 2000 heating steps, 5000 cooling steps, and a final cooling target temperature of 300 K.

Fig. S1. RD cells were infected with EV-D68 (IL/14-18952) at 0.1 MOI. Cells were treated with 5 µM of indicated compound and images were captured at 72 hours post infection Representative images (A) are shown.

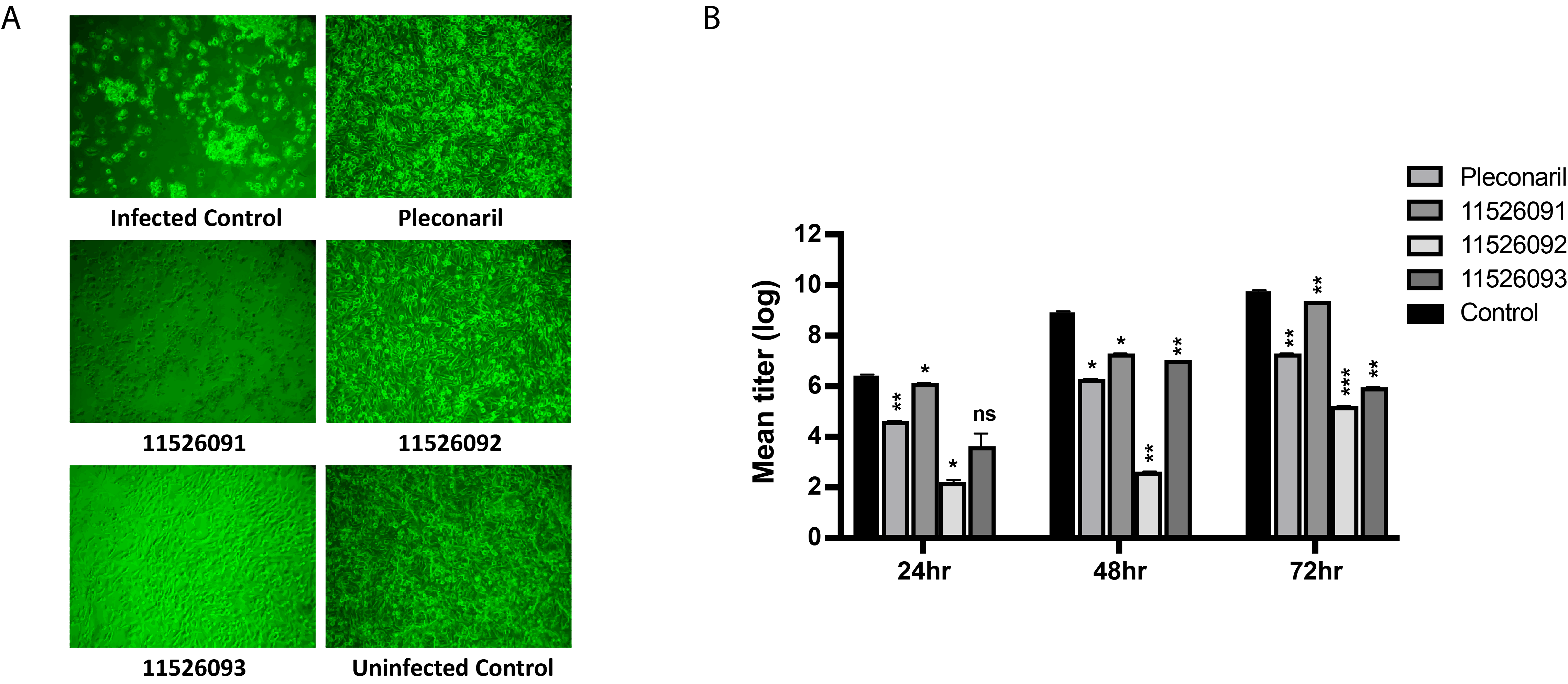

Fig. S2. Structural (A) and sequence alignment (B) of multiple analogous capsid structures of select enteroviruses. EV-D68 (yellow; 4WM7), EV-A71 (light grey; 4CEW), Coxsackievirus A10 (cyan; 6SNB), poliovirus type 3 (green; 5O5B), rhinovirus 16 (dark grey; 1AYM), Coxsackievirus B3 (blue; 4GB3).

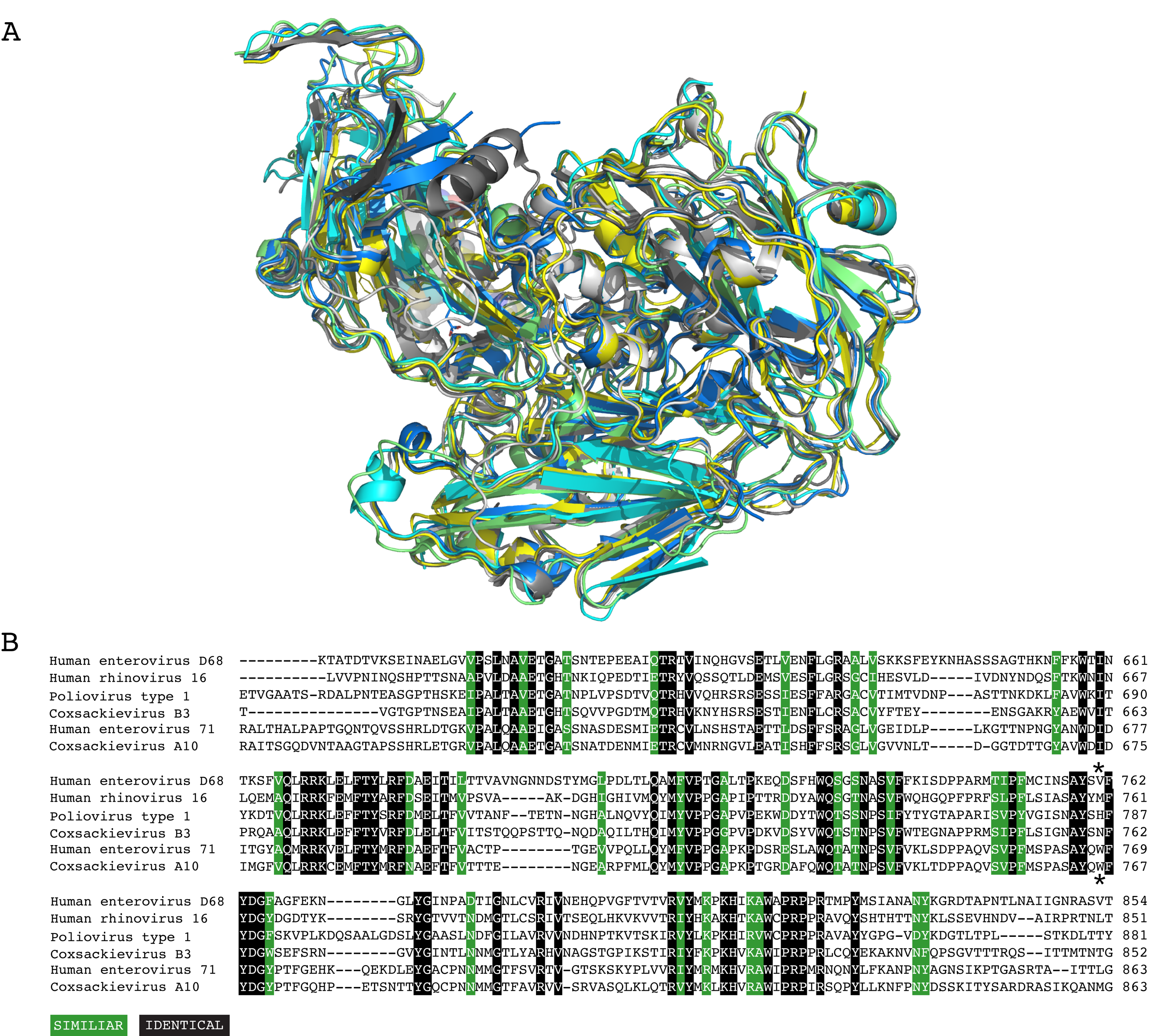

**Fig. S3.** MTD study with 2-week (A-B) and 10-day-old (C-D) mice treated with 11526092 by oral gavage and IP administration, respectively. Survival is not significantly different from the control groups for IP or oral administration, but there was a minor, but statistically significant, % weight gain found in the 10 (Adjusted p=0.0185) and 30 (Adjusted p=0.0148) mg/kg/day oral-administration groups over vehicle. ANOVA (Brown-Forsythe and Welch) followed by a Dunnett’s T3 multiple comparisons test was performed in Graphpad Prism 9.2. Error bars represent the SEM.

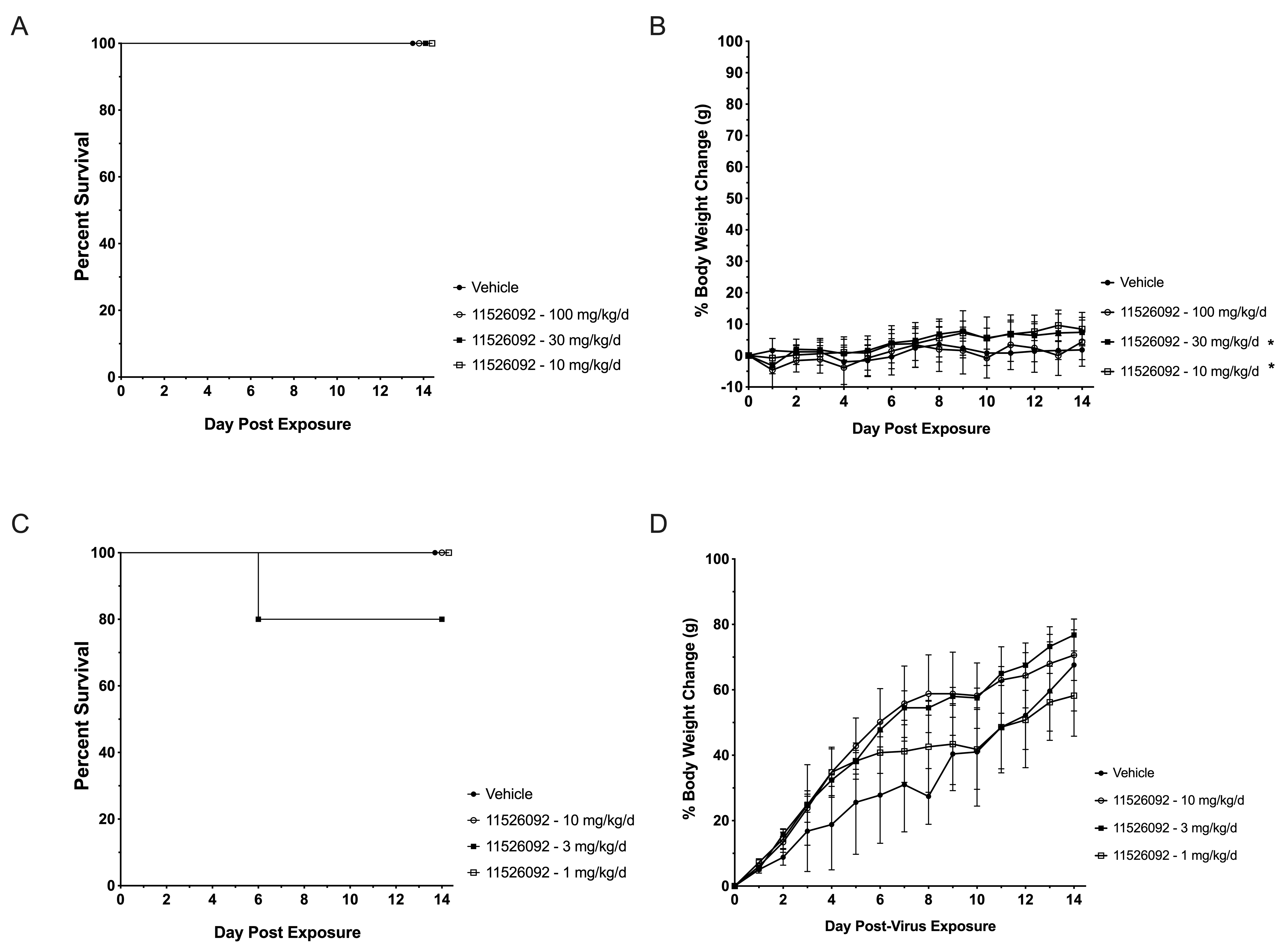

**Fig. S4.** AFM respiratory model weight loss post virus exposure. The mean % body weight change of each maEV-D68-exposed, treated groups as compared to the placebo control group. Statistically significant % weight loss mitigation versus placebo was found in the virus-challenged 11526092 100 mg/kg/d group only. Statistical analysis determined by an ANOVA (Brown-Forsythe and Welch) followed by a Dunnett’s T3 multiple comparisons tests were performed in Graphpad Prism 9.2. Error bars represent the SEM.

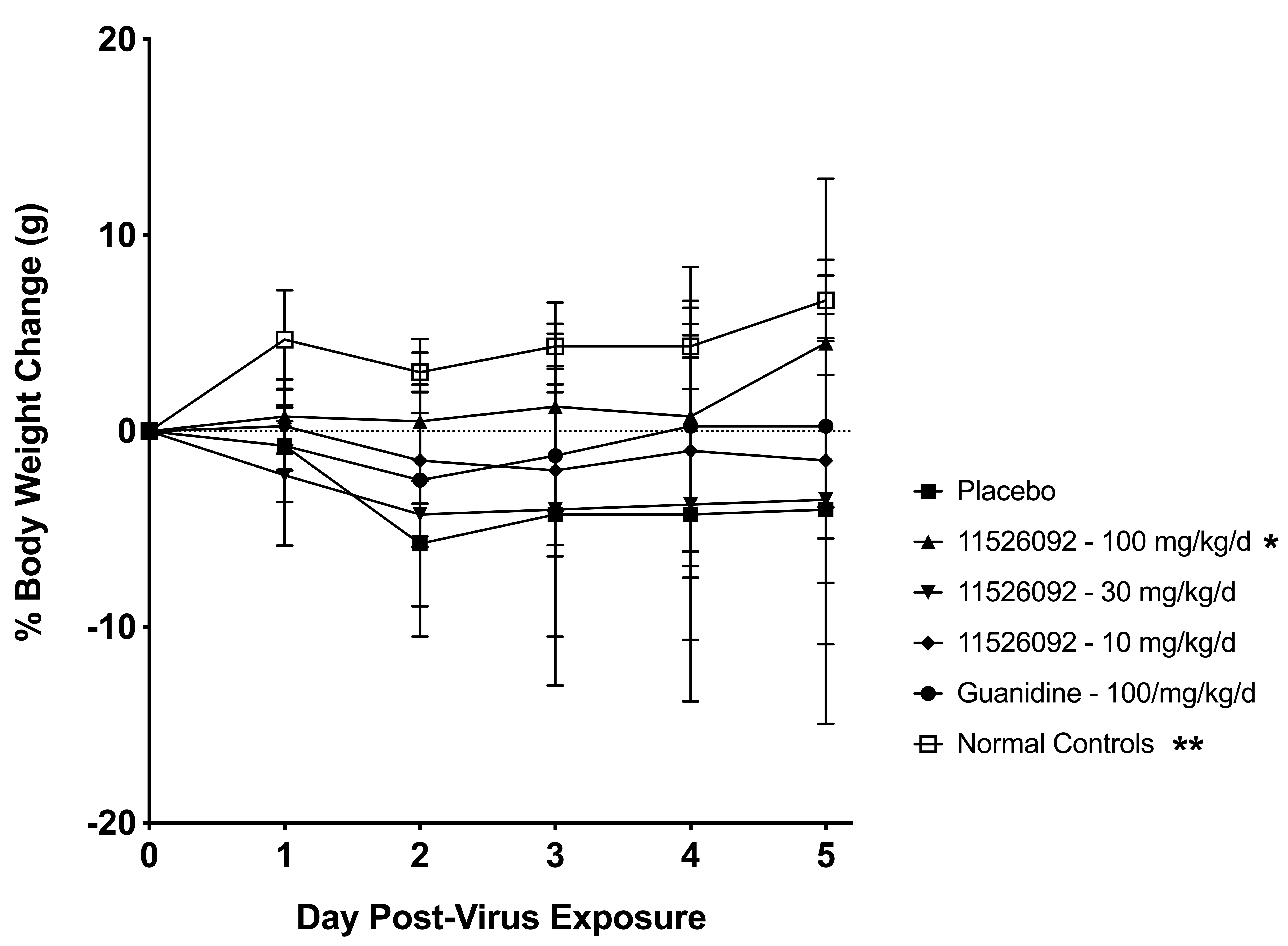

**Fig. S5.** Select cytokine and interleukins levels following ma-EV-D68 challenge in the AFM respiratory mouse model. Each cytokine/interleukin level was compared to the placebo group for statistical significance. Ordinary two-way ANOVA followed by a Dunnett’s T3 multiple comparisons test was performed in Graphpad Prism 9.2 for macOS. Error bars represent the SEM.

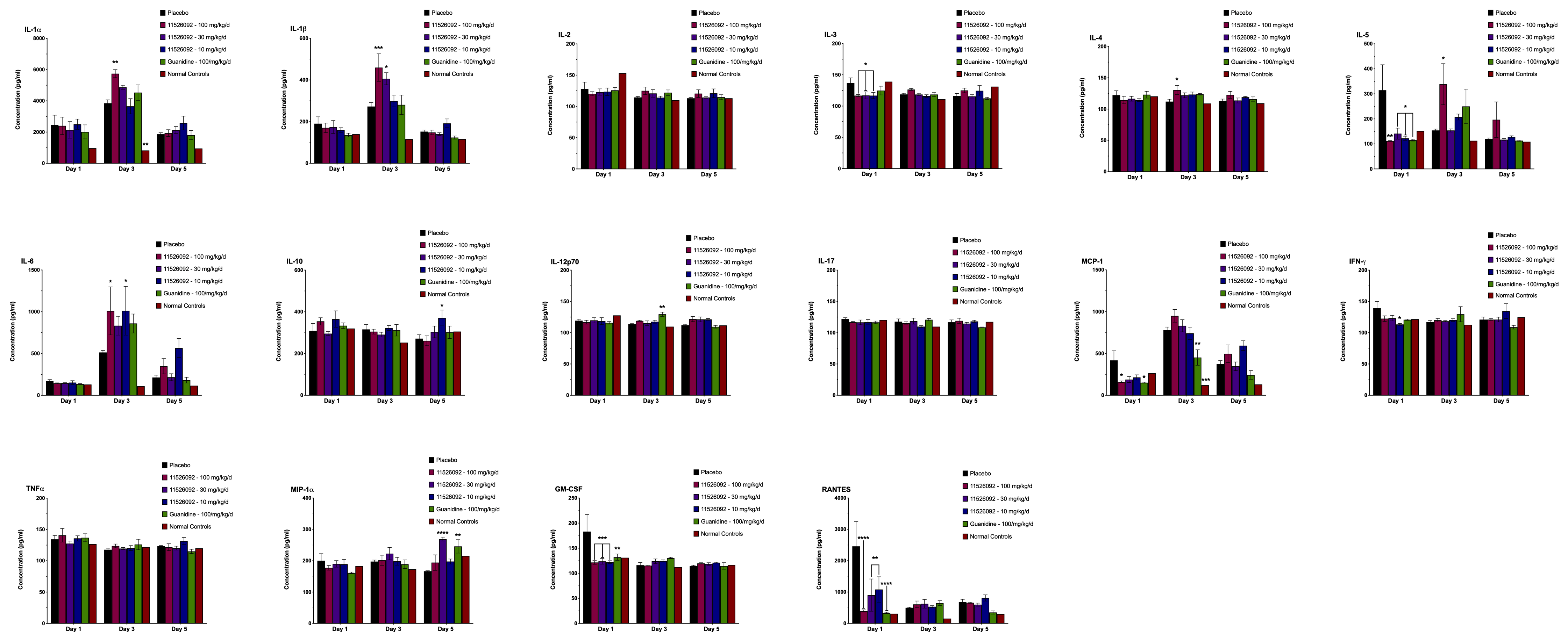

**Fig. S6.** AFM neurological data: In both the pre- and post-challenge treated groups only the positive control Guanidine-treated group showed a significant increase in survival. Neurological scores are shown for days 1-11 and 1-4 for mice treated pre- and post-challenge, respectively. None of the 1526092-treated mice showed a reduction in neurological scores. Guanidine, used as positive control, reduced neurological scores on day 1 with pre-challenge administration and day 3 post-challenge administration. Only the Guanidine treated group survived past day 4 in the post-challenge treated groups, making a comparison impossible after day 4. (*P<0.05 compared to placebo-treated mice). Viremia levels also only showed a statically significant reduction in the control groups. Survival curves were assessed for differences using a log-rank (Mantel-Cox) test. Blood titer and neurological score differences were determined using an ordinary one-way ANOVA followed by a Dunnett’s T3 multiple comparisons test. All analyses were performed in Graphpad Prism 9.1.1. Error bars represent the SEM.

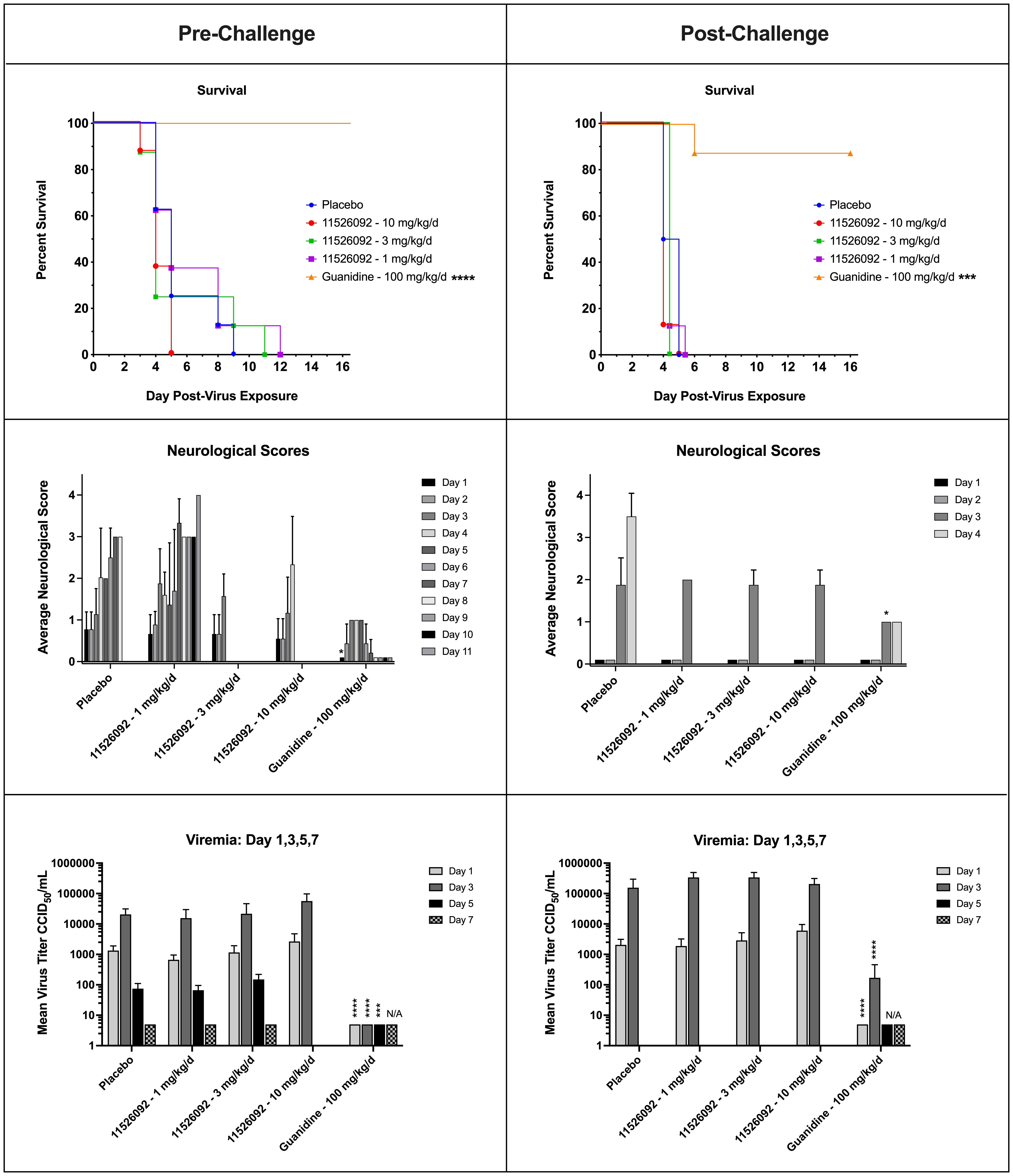

**Fig. S7.** Percent body weight change of treated mice in AFM neurological models. (A) For the pre-challenge treatment study, the % body weight change was statistically different from the placebo group for 11526092 (3 mg/kg), guanidine-treated (positive control) and normal, uninfected control groups. For treatment post-challenge, (B) none of the treated groups showed a significant improvement in body weight gain as compared to the placebo control. % Weight change differences were determined using an ANOVA (Brown-Forsythe and Welch) followed by a Dunnett’s T3 multiple comparisons test. All analyses were performed in Graphpad Prism 9.2. It is noted that due to the stark difference in survival rates between groups these differences are unlikely to be reliable as all the placebo-treated animals died or were euthanized by day 4 and 8 for the pre- and post-treated groups, respectively. Error bars represent the SEM.

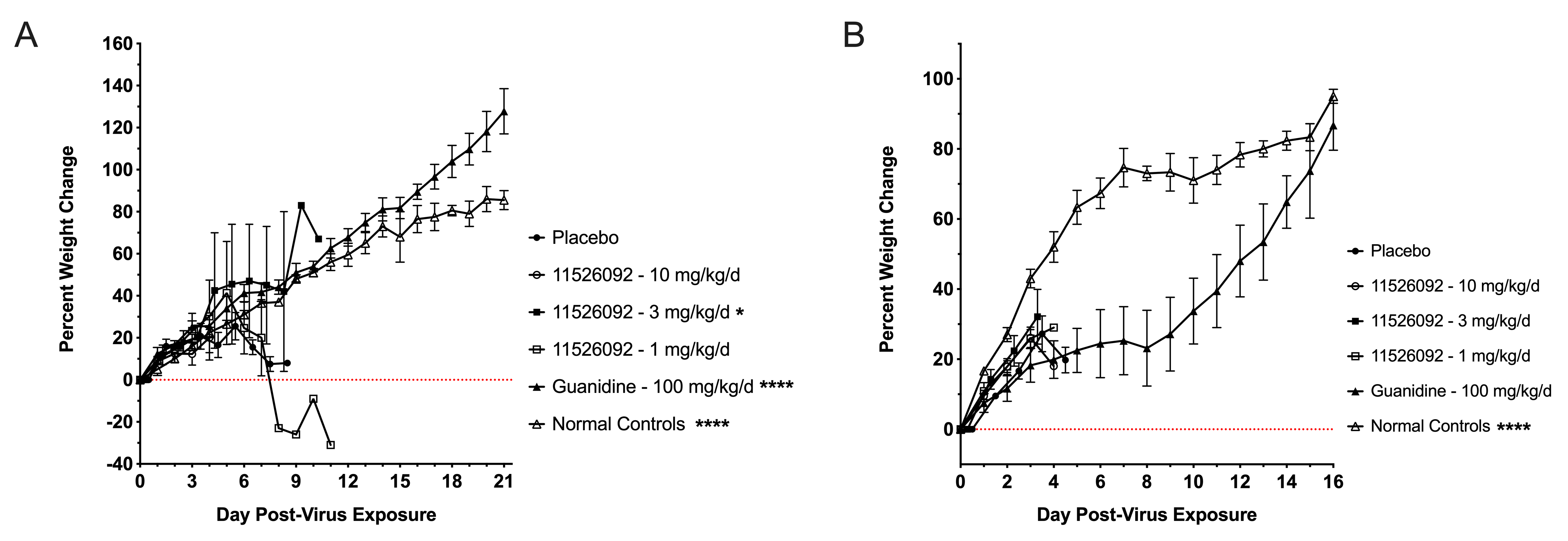

Fig. S8. Docking of 11526092 in the previously published EV-D68 (Fermon) capsid structure. (A) Pleconaril (salmon) in the EV-D68 capsid structure PDB: 4WM7 (yellow; pleconaril ligand in crystal) represented as a cartoon with a highlighted juxtaposed hydrophobic residue (VAL-195) shown as spheres (representing the Van der Waals radius). (B) The top scoring pose of 11526092 (orange) using CDOCKER (CDOCKER Energy 44.7566) suggests that this molecule picks up an additional hydrophobic interaction with VAL-195 (VP1) not found in the previously reported pleconaril-EV-D68 (Fermon) capsid crystal structure. This is substantiated in the cryo-EM structure of EV68 (MO strain) complexed with 11526092 (C). *These data* suggest this may be a potential reason for the increased potency of 11526092 versus pleconaril against EV-D68 *in vitro*. As a reference, the top scoring pleconaril pose had a CDOCKER Energy of 41.3514 with a RMSD of 0.681 Å from the crystal structure. (D) Illustrating how 11526092 may pick up hydrophobic interactions versus other analogs without the methyl group, which suggests a rationale for their inhibition differences.

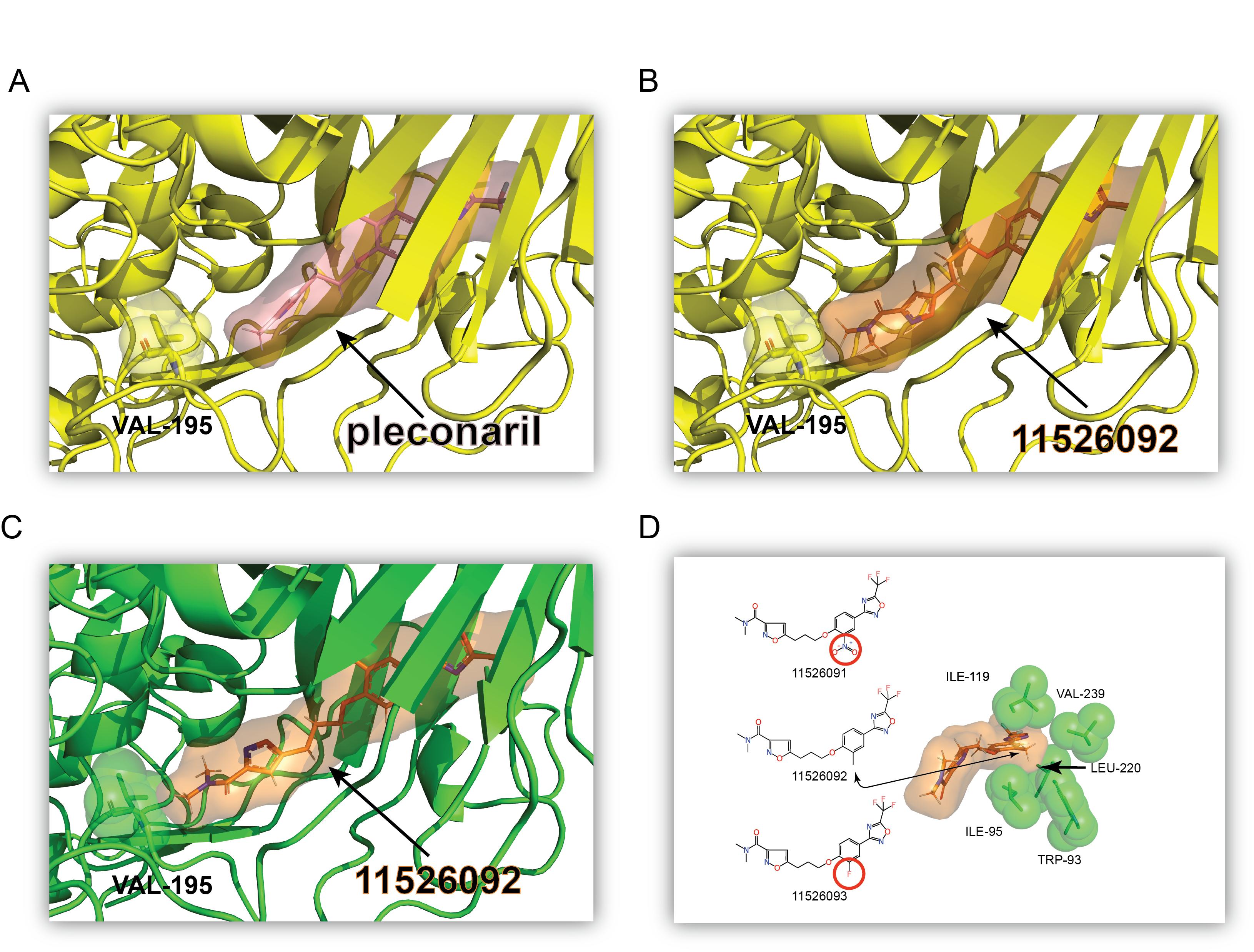
**Fig. S9: Interface comparison between EV-D68-pleconaril (Fermon), EV-D68-11526092 (MO) and EV-D68-pleconaril (MO).** (A) Pleconaril and interfacing residues (Fermon; PDB: 4wm7). (B) 11526092 and interfacing residues. (C) Pleconaril and interfacing residues (MO). Residues at the interface are highly conserved. Differences between (A) (B) and (B) (C) are highlighted in red and blue.

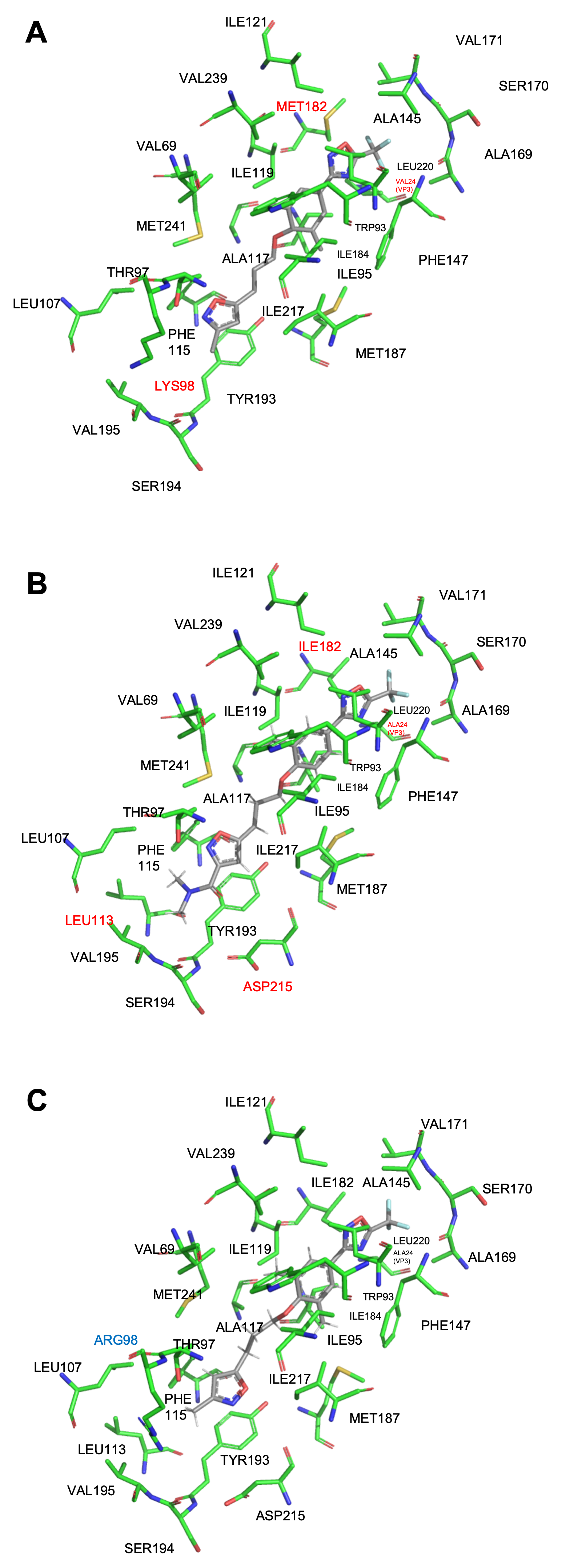

**Fig. S10: Comparison of the conformational changes induced by pleconaril and 11526092.** (A) Based on the structure of EV-D68 Fermon and pleconaril (PDB: 4wm7) (cyan) and the structure of EV-D68 Fermon (PDB: 4wm8) (transparent blue), pleconaril triggers an inward (cyan arrow) movement of a residue on the GH loop of VP1. (B) (C) The VP1 proteins of EV-D68MO-11526902 (blue) and EV-D68MO-Pleconaril (green) are compared with the VP1 of EV-D68 MO (transparent purple). Both viral-inhibitor complexes show outward (blue and green arrows) movements of VP1 GH and EF loops.

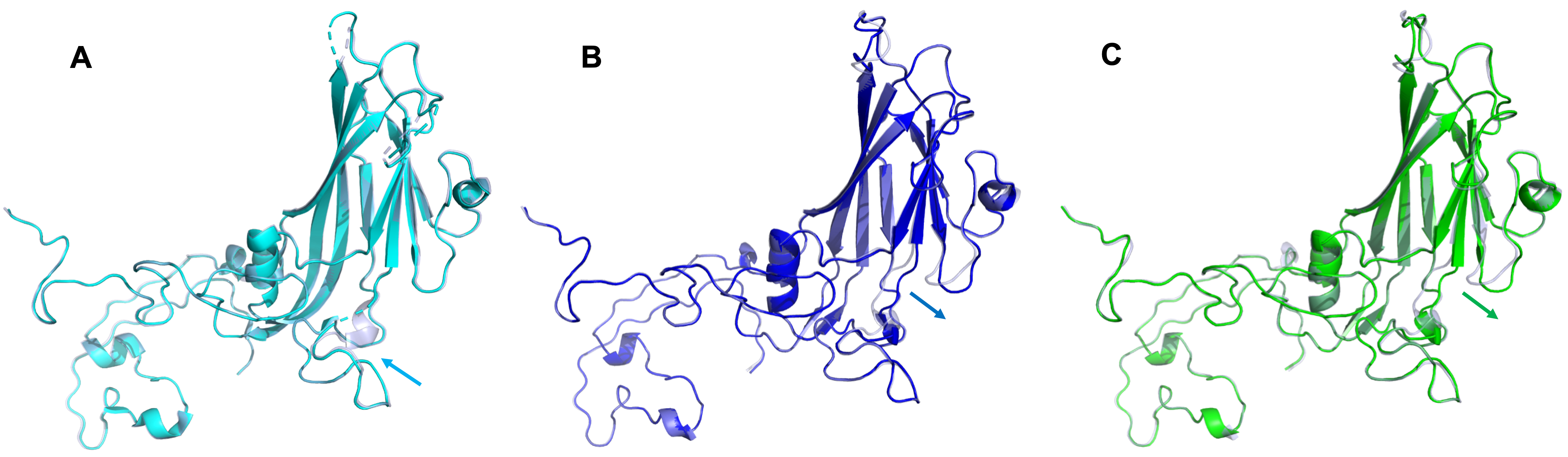

**Fig S11:** Superimposition of EV-D68 MO in complex with pleconaril (gold) and (A) EV-D68MO native virion (green), (B) EV-D68MO in complex with 11526092 (blue), and (C) EV-D68 Fermon in complex with pleconaril (PDB:4wm7) (pink). The essential region on VP1 GH loop is highlighted using black arrows. Unlike the complex of EV-D68 Fermon with pleconaril, for the EV-D68 MO strain, pleconaril binds and pushes the pocket roof outwards, showing a similar movement as the complex of EV-D68 MO and 11526092.

A	B	C

**Fig S12:** A close-up view of EV-D68 MO in complex of pleconaril (gold) and EV-D68 Fermon in complex with pleconaril (PDB:4wm7) (pink). In contrast to the inward movement of VP1 residue 211 (red circle), which possibly blocks the pocket entrance, observed in the EV-D68 Fermon and pleconaril complexes, an outward movement of the loop that forms the pocket roof is observed with EV-D68 MO.

**Fig. S13.** CYP3A4 activation at high concentrations of 11526092 and pleconaril. Error bars represent the SEM (n=2). For 11526092 the first major enhancement was shown at 0.780 µM, with metabolism of testosterone at 173.5 ± 42.17% (±SEM) of control.

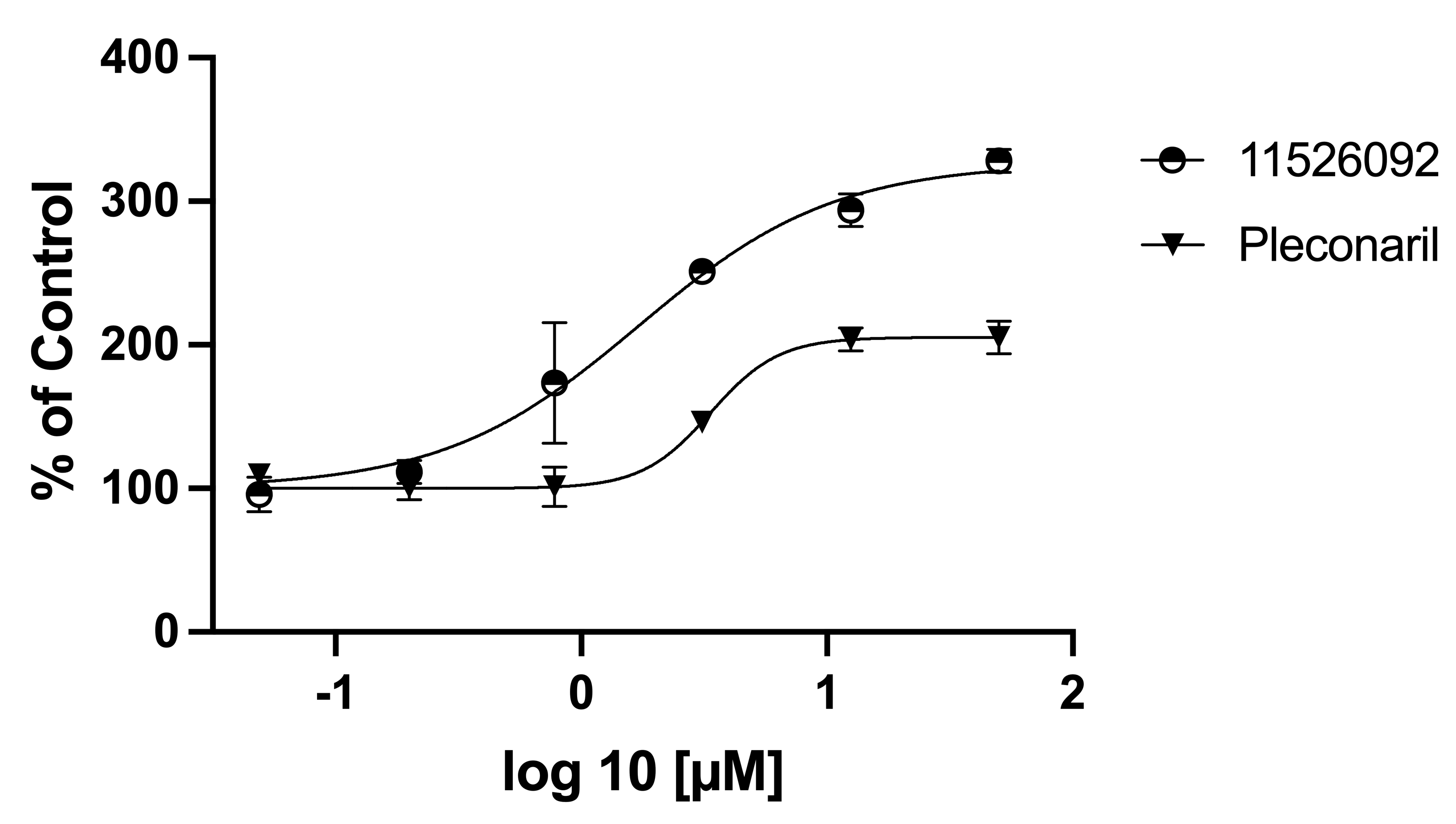

**Fig. S14: Density of the VP1 hydrophobic pocket region.** The residues and densities at VP1 pocket regions of (A) EV-D68 MO alone, (B) EV-D68 MO and 11526091 and (C) EV-D68 MO and 11526093 are shown as sticks and meshes (contour levels around 0.8). Similar as the pocket region of the native virion, there are no obvious densities observed for ligand fitting in the pocket regions for 11526091 and 11526093. Other than empty pockets, there are no significant conformational changes observed. Overall, the structures show no binding or very low occupancy for samples of EV-D68 MO incubated with 11526091 and 11526093**.**

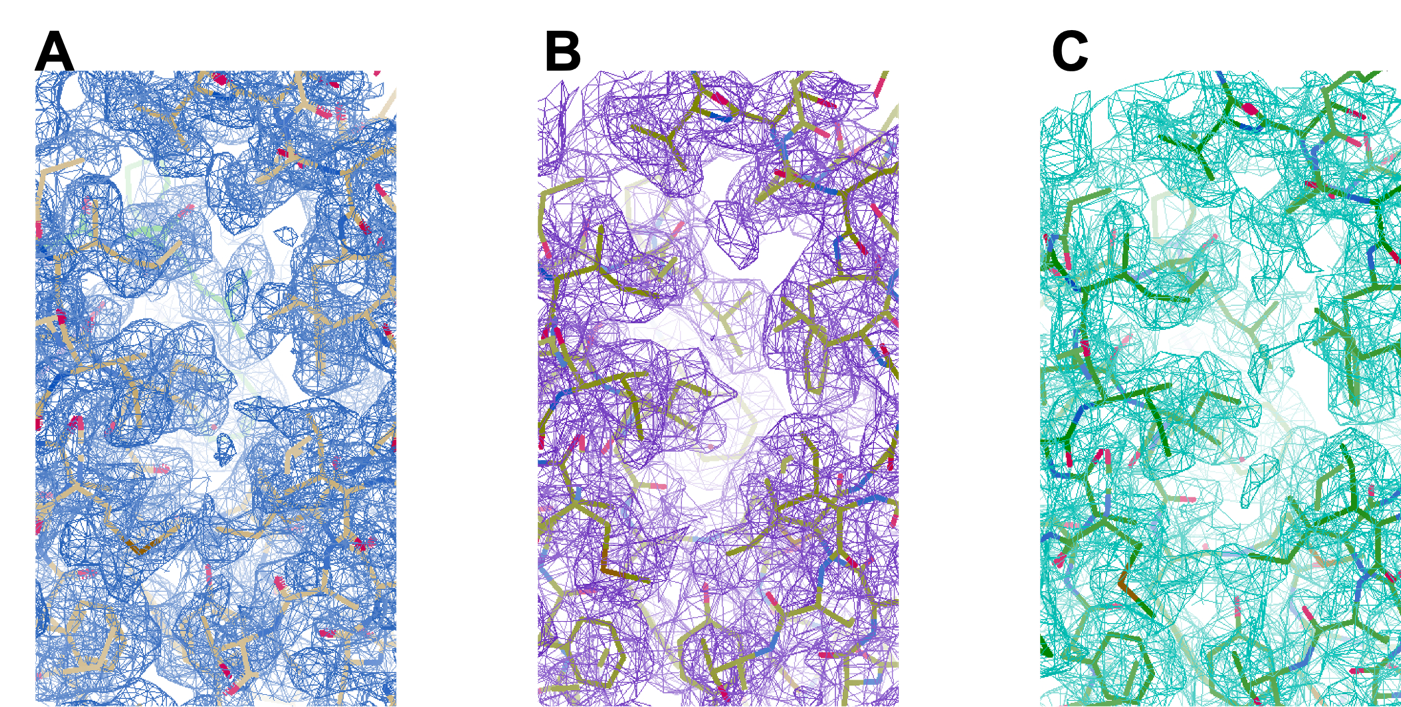

**Fig. S15: Density for Pleconaril and 11526092 in EV-D68 MO VP1 hydrophobic pocket regions.** Residues lining the pocket are based on Liu et al, 2015 (*17*), and are shown in green. The densities of (A) pleconaril and (B) 11526092 are colored in red and blue, respectively.

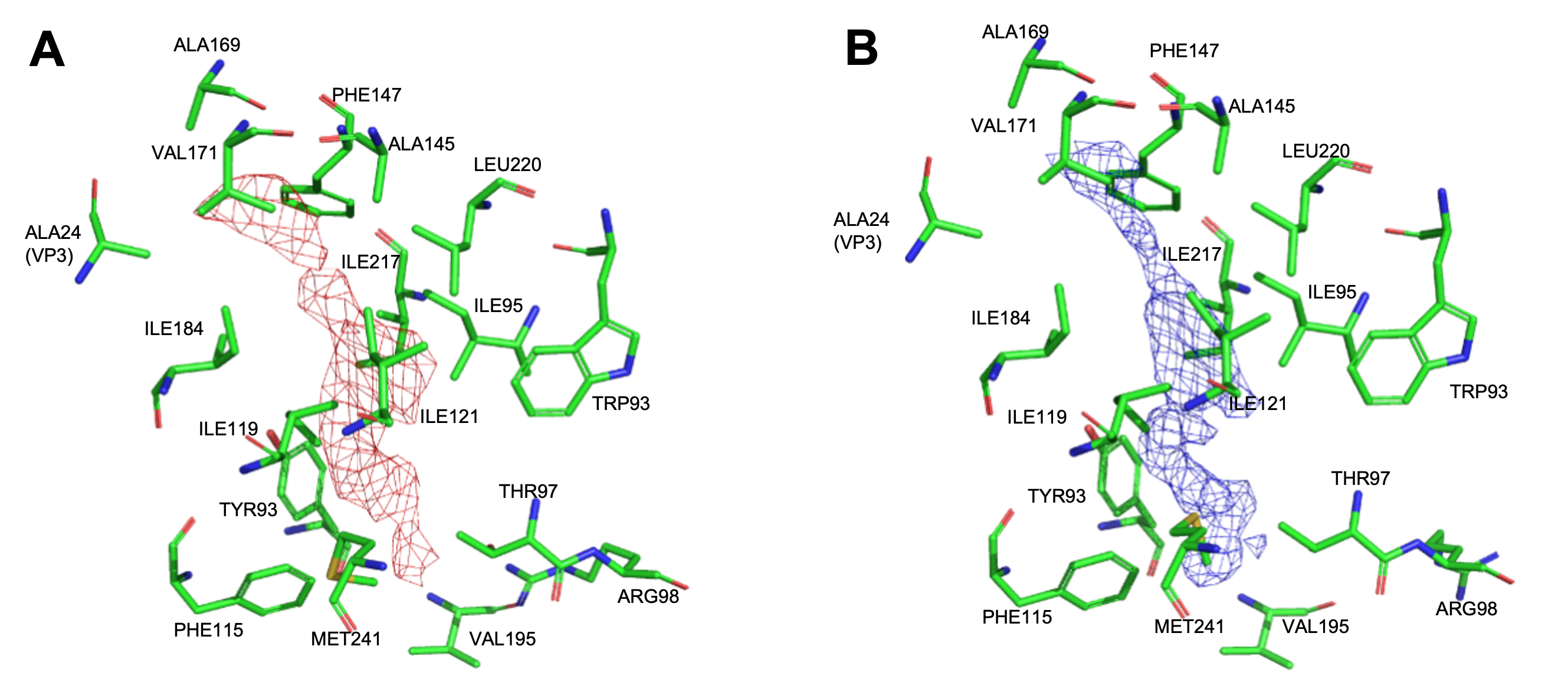

**Fig. S16:** Example of typical map densities at 2 Å resolution. VP1, VP2, VP3 and VP4 residues are shown in sticks and colored in blue, green, red, and magenta, respectively.

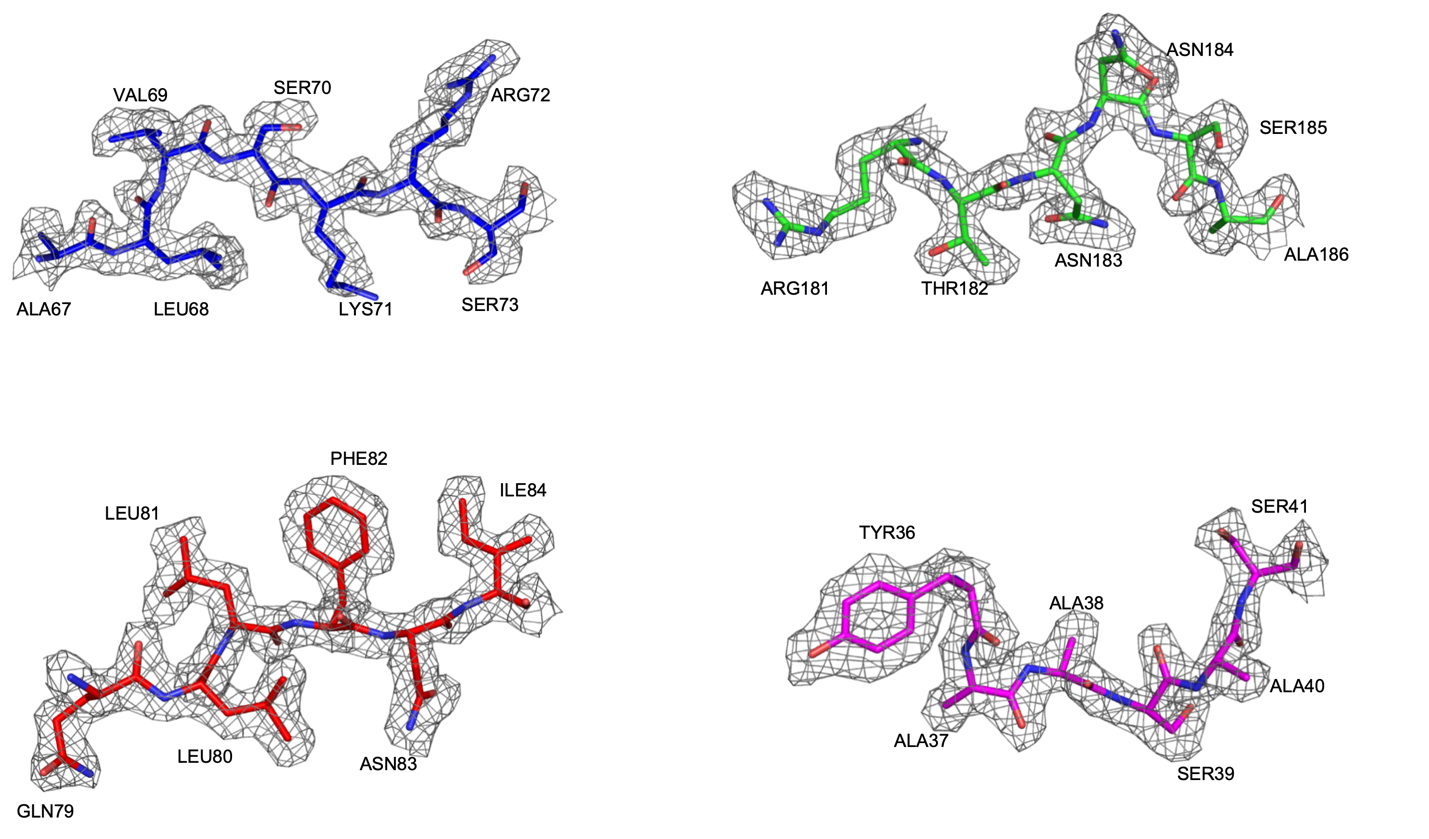

**Fig. S17:** Estimates of map resolution of EV-D68 MO natural virion (purple), EV-D68 MO in complex with 11526092 (red), EV-D68 MO in complex with pleconaril (blue), EV-D68 MO in complex with 11526091 (black) and EV-D68 MO in complex with 11526093 (green) are based on the gold-standard Fourier shell correlation cutoff of 0.143 (magenta).

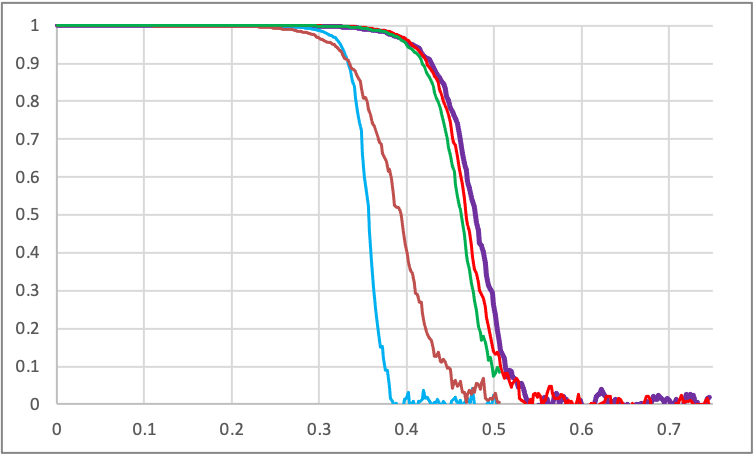

FSC=0.143

1/Resolution(Å^-1^)

Correlation Coefficient

|  | frequency |
| --- | --- |
| 0.000993057 | 1 |
| 0.00297917 | 1 |
| 0.00496528 | 1 |
| 0.0069514 | 1 |
| 0.00893751 | 1 |
| 0.0109236 | 0.999999 |
| 0.0129097 | 1 |
| 0.0148958 | 1 |
| 0.016882 | 1 |
| 0.0188681 | 0.999999 |
| 0.0208542 | 0.999999 |
| 0.0228403 | 0.999996 |
| 0.0248264 | 0.999997 |
| 0.0268125 | 0.999998 |
| 0.0287986 | 0.999994 |
| 0.0307848 | 0.999997 |
| 0.0327709 | 0.999995 |
| 0.034757 | 0.999995 |
| 0.0367431 | 0.999995 |
| 0.0387292 | 0.999994 |
| 0.0407153 | 0.999993 |
| 0.0427014 | 0.999988 |
| 0.0446875 | 0.999992 |
| 0.0466737 | 0.999984 |
| 0.0486598 | 0.999986 |
| 0.0506459 | 0.999982 |
| 0.052632 | 0.999975 |
| 0.0546181 | 0.999965 |
| 0.0566042 | 0.999969 |
| 0.0585903 | 0.999972 |
| 0.0605764 | 0.999963 |
| 0.0625626 | 0.999945 |
| 0.0645487 | 0.999964 |
| 0.0665348 | 0.99996 |
| 0.0685209 | 0.999942 |
| 0.070507 | 0.999947 |
| 0.0724931 | 0.999956 |
| 0.0744792 | 0.999956 |
| 0.0764654 | 0.999964 |
| 0.0784515 | 0.999959 |
| 0.0804376 | 0.999939 |
| 0.0824237 | 0.999955 |
| 0.0844098 | 0.999953 |
| 0.0863959 | 0.999947 |
| 0.088382 | 0.999952 |
| 0.0903681 | 0.999959 |
| 0.0923543 | 0.999949 |
| 0.0943404 | 0.999949 |
| 0.0963265 | 0.999953 |
| 0.0983126 | 0.999959 |
| 0.100299 | 0.999962 |
| 0.102285 | 0.999958 |
| 0.104271 | 0.999961 |
| 0.106257 | 0.999961 |
| 0.108243 | 0.999947 |
| 0.110229 | 0.999926 |
| 0.112215 | 0.999934 |
| 0.114202 | 0.999896 |
| 0.116188 | 0.99991 |
| 0.118174 | 0.999882 |
| 0.12016 | 0.999911 |
| 0.122146 | 0.999878 |
| 0.124132 | 0.999867 |
| 0.126118 | 0.999898 |
| 0.128104 | 0.999885 |
| 0.13009 | 0.999858 |
| 0.132077 | 0.999867 |
| 0.134063 | 0.999844 |
| 0.136049 | 0.999855 |
| 0.138035 | 0.999812 |
| 0.140021 | 0.99982 |
| 0.142007 | 0.999821 |
| 0.143993 | 0.999849 |
| 0.145979 | 0.999854 |
| 0.147965 | 0.999848 |
| 0.149952 | 0.999869 |
| 0.151938 | 0.999855 |
| 0.153924 | 0.999839 |
| 0.15591 | 0.999845 |
| 0.157896 | 0.999827 |
| 0.159882 | 0.999815 |
| 0.161868 | 0.999812 |
| 0.163854 | 0.99982 |
| 0.16584 | 0.999819 |
| 0.167827 | 0.999828 |
| 0.169813 | 0.999843 |
| 0.171799 | 0.999825 |
| 0.173785 | 0.999813 |
| 0.175771 | 0.999807 |
| 0.177757 | 0.999786 |
| 0.179743 | 0.99983 |
| 0.181729 | 0.99983 |
| 0.183715 | 0.999852 |
| 0.185702 | 0.999854 |
| 0.187688 | 0.99984 |
| 0.189674 | 0.999843 |
| 0.19166 | 0.999811 |
| 0.193646 | 0.999831 |
| 0.195632 | 0.999826 |
| 0.197618 | 0.999837 |
| 0.199604 | 0.999844 |
| 0.20159 | 0.999824 |
| 0.203577 | 0.999876 |
| 0.205563 | 0.999845 |
| 0.207549 | 0.999851 |
| 0.209535 | 0.999853 |
| 0.211521 | 0.999834 |
| 0.213507 | 0.999833 |
| 0.215493 | 0.999814 |
| 0.217479 | 0.999823 |
| 0.219465 | 0.999803 |
| 0.221452 | 0.999797 |
| 0.223438 | 0.999798 |
| 0.225424 | 0.999798 |
| 0.22741 | 0.999819 |
| 0.229396 | 0.999827 |
| 0.231382 | 0.999808 |
| 0.233368 | 0.999816 |
| 0.235354 | 0.999782 |
| 0.237341 | 0.999749 |
| 0.239327 | 0.999769 |
| 0.241313 | 0.999699 |
| 0.243299 | 0.999696 |
| 0.245285 | 0.999718 |
| 0.247271 | 0.999636 |
| 0.249257 | 0.999627 |
| 0.251243 | 0.999601 |
| 0.253229 | 0.999555 |
| 0.255216 | 0.999522 |
| 0.257202 | 0.999513 |
| 0.259188 | 0.999519 |
| 0.261174 | 0.999471 |
| 0.26316 | 0.99947 |
| 0.265146 | 0.999455 |
| 0.267132 | 0.999429 |
| 0.269118 | 0.999411 |
| 0.271104 | 0.999358 |
| 0.273091 | 0.999284 |
| 0.275077 | 0.999253 |
| 0.277063 | 0.99923 |
| 0.279049 | 0.999156 |
| 0.281035 | 0.999034 |
| 0.283021 | 0.998996 |
| 0.285007 | 0.999037 |
| 0.286993 | 0.998938 |
| 0.288979 | 0.998842 |
| 0.290966 | 0.998803 |
| 0.292952 | 0.998785 |
| 0.294938 | 0.998713 |
| 0.296924 | 0.998549 |
| 0.29891 | 0.998592 |
| 0.300896 | 0.998507 |
| 0.302882 | 0.998332 |
| 0.304868 | 0.998132 |
| 0.306854 | 0.997985 |
| 0.308841 | 0.998044 |
| 0.310827 | 0.997869 |
| 0.312813 | 0.99775 |
| 0.314799 | 0.997589 |
| 0.316785 | 0.997572 |
| 0.318771 | 0.997312 |
| 0.320757 | 0.997163 |
| 0.322743 | 0.996822 |
| 0.324729 | 0.996611 |
| 0.326716 | 0.996412 |
| 0.328702 | 0.99619 |
| 0.330688 | 0.996049 |
| 0.332674 | 0.995728 |
| 0.33466 | 0.995512 |
| 0.336646 | 0.995149 |
| 0.338632 | 0.994735 |
| 0.340618 | 0.994337 |
| 0.342605 | 0.994125 |
| 0.344591 | 0.993429 |
| 0.346577 | 0.992791 |
| 0.348563 | 0.991738 |
| 0.350549 | 0.990757 |
| 0.352535 | 0.990236 |
| 0.354521 | 0.989585 |
| 0.356507 | 0.989122 |
| 0.358493 | 0.988838 |
| 0.36048 | 0.988081 |
| 0.362466 | 0.987692 |
| 0.364452 | 0.986697 |
| 0.366438 | 0.986058 |
| 0.368424 | 0.985482 |
| 0.37041 | 0.985004 |
| 0.372396 | 0.983741 |
| 0.374382 | 0.983524 |
| 0.376368 | 0.981717 |
| 0.378355 | 0.979111 |
| 0.380341 | 0.97724 |
| 0.382327 | 0.975558 |
| 0.384313 | 0.972949 |
| 0.386299 | 0.972261 |
| 0.388285 | 0.970842 |
| 0.390271 | 0.970053 |
| 0.392257 | 0.968275 |
| 0.394243 | 0.966884 |
| 0.39623 | 0.965414 |

|  | frequency |
| --- | --- |
| 0.000996492 | 1 |
| 0.00298948 | 1 |
| 0.00498246 | 1 |
| 0.00697545 | 1 |
| 0.00896843 | 1 |
| 0.0109614 | 0.999999 |
| 0.0129544 | 0.999998 |
| 0.0149474 | 0.999999 |
| 0.0169404 | 0.999995 |
| 0.0189334 | 0.999994 |
| 0.0209263 | 0.999987 |
| 0.0229193 | 0.999987 |
| 0.0249123 | 0.999982 |
| 0.0269053 | 0.999965 |
| 0.0288983 | 0.999936 |
| 0.0308913 | 0.999961 |
| 0.0328842 | 0.999956 |
| 0.0348772 | 0.999969 |
| 0.0368702 | 0.999981 |
| 0.0388632 | 0.999961 |
| 0.0408562 | 0.999945 |
| 0.0428492 | 0.999939 |
| 0.0448422 | 0.999936 |
| 0.0468351 | 0.999911 |
| 0.0488281 | 0.999898 |
| 0.0508211 | 0.999842 |
| 0.0528141 | 0.999872 |
| 0.0548071 | 0.999851 |
| 0.0568001 | 0.999836 |
| 0.058793 | 0.999846 |
| 0.060786 | 0.999869 |
| 0.062779 | 0.999846 |
| 0.064772 | 0.999884 |
| 0.066765 | 0.999861 |
| 0.068758 | 0.999848 |
| 0.070751 | 0.999865 |
| 0.0727439 | 0.999888 |
| 0.0747369 | 0.999877 |
| 0.0767299 | 0.99988 |
| 0.0787229 | 0.99987 |
| 0.0807159 | 0.999867 |
| 0.0827089 | 0.999884 |
| 0.0847018 | 0.99988 |
| 0.0866948 | 0.999854 |
| 0.0886878 | 0.99987 |
| 0.0906808 | 0.999897 |
| 0.0926738 | 0.999897 |
| 0.0946668 | 0.999882 |
| 0.0966598 | 0.999867 |
| 0.0986527 | 0.999887 |
| 0.100646 | 0.999886 |
| 0.102639 | 0.999878 |
| 0.104632 | 0.999893 |
| 0.106625 | 0.999851 |
| 0.108618 | 0.999828 |
| 0.110611 | 0.999828 |
| 0.112604 | 0.999813 |
| 0.114597 | 0.999758 |
| 0.11659 | 0.999706 |
| 0.118583 | 0.999727 |
| 0.120576 | 0.999757 |
| 0.122569 | 0.999723 |
| 0.124562 | 0.999698 |
| 0.126555 | 0.999745 |
| 0.128548 | 0.999723 |
| 0.13054 | 0.99968 |
| 0.132533 | 0.999593 |
| 0.134526 | 0.999581 |
| 0.136519 | 0.99957 |
| 0.138512 | 0.999561 |
| 0.140505 | 0.999527 |
| 0.142498 | 0.999541 |
| 0.144491 | 0.999529 |
| 0.146484 | 0.999526 |
| 0.148477 | 0.999532 |
| 0.15047 | 0.999553 |
| 0.152463 | 0.999508 |
| 0.154456 | 0.999466 |
| 0.156449 | 0.999495 |
| 0.158442 | 0.999467 |
| 0.160435 | 0.999419 |
| 0.162428 | 0.999387 |
| 0.164421 | 0.999359 |
| 0.166414 | 0.99948 |
| 0.168407 | 0.999498 |
| 0.1704 | 0.999514 |
| 0.172393 | 0.999487 |
| 0.174386 | 0.999456 |
| 0.176379 | 0.999423 |
| 0.178372 | 0.999399 |
| 0.180365 | 0.999467 |
| 0.182358 | 0.999417 |
| 0.184351 | 0.999419 |
| 0.186344 | 0.999399 |
| 0.188337 | 0.999398 |
| 0.19033 | 0.99941 |
| 0.192323 | 0.999393 |
| 0.194316 | 0.999448 |
| 0.196309 | 0.999472 |
| 0.198302 | 0.999492 |
| 0.200295 | 0.999477 |
| 0.202288 | 0.99945 |
| 0.204281 | 0.999482 |
| 0.206274 | 0.999477 |
| 0.208267 | 0.99951 |
| 0.21026 | 0.999487 |
| 0.212253 | 0.999488 |
| 0.214246 | 0.999471 |
| 0.216239 | 0.999428 |
| 0.218232 | 0.999442 |
| 0.220225 | 0.99942 |
| 0.222218 | 0.999329 |
| 0.224211 | 0.999299 |
| 0.226204 | 0.999279 |
| 0.228197 | 0.999252 |
| 0.23019 | 0.999249 |
| 0.232183 | 0.999236 |
| 0.234176 | 0.999203 |
| 0.236169 | 0.999091 |
| 0.238162 | 0.998933 |
| 0.240155 | 0.998905 |
| 0.242148 | 0.998808 |
| 0.244141 | 0.998852 |
| 0.246134 | 0.998789 |
| 0.248127 | 0.998563 |
| 0.25012 | 0.998555 |
| 0.252113 | 0.998433 |
| 0.254106 | 0.998321 |
| 0.256099 | 0.998119 |
| 0.258092 | 0.998106 |
| 0.260085 | 0.997654 |
| 0.262077 | 0.997552 |
| 0.26407 | 0.997362 |
| 0.266063 | 0.997051 |
| 0.268056 | 0.996884 |
| 0.270049 | 0.996751 |
| 0.272042 | 0.996163 |
| 0.274035 | 0.996073 |
| 0.276028 | 0.995976 |
| 0.278021 | 0.995408 |
| 0.280014 | 0.994646 |
| 0.282007 | 0.994313 |
| 0.284 | 0.99386 |
| 0.285993 | 0.993471 |
| 0.287986 | 0.992814 |
| 0.289979 | 0.991629 |
| 0.291972 | 0.990951 |
| 0.293965 | 0.990526 |
| 0.295958 | 0.98984 |
| 0.297951 | 0.988358 |
| 0.299944 | 0.987009 |
| 0.301937 | 0.98443 |
| 0.30393 | 0.983518 |
| 0.305923 | 0.98125 |
| 0.307916 | 0.978115 |
| 0.309909 | 0.97745 |
| 0.311902 | 0.974671 |
| 0.313895 | 0.973063 |
| 0.315888 | 0.970053 |
| 0.317881 | 0.96823 |
| 0.319874 | 0.963565 |
| 0.321867 | 0.957753 |
| 0.32386 | 0.950924 |
| 0.325853 | 0.945631 |
| 0.327846 | 0.933588 |
| 0.329839 | 0.921916 |
| 0.331832 | 0.908561 |
| 0.333825 | 0.896746 |
| 0.335818 | 0.880452 |
| 0.337811 | 0.853838 |
| 0.339804 | 0.838547 |
| 0.341797 | 0.810688 |
| 0.34379 | 0.773601 |
| 0.345783 | 0.74484 |
| 0.347776 | 0.722732 |
| 0.349769 | 0.66012 |
| 0.351762 | 0.600118 |
| 0.353755 | 0.561015 |
| 0.355748 | 0.521215 |
| 0.357741 | 0.456907 |
| 0.359734 | 0.380477 |
| 0.361727 | 0.310584 |
| 0.36372 | 0.255587 |
| 0.365713 | 0.230777 |
| 0.367706 | 0.19102 |
| 0.369699 | 0.150607 |
| 0.371692 | 0.151758 |
| 0.373685 | 0.121084 |
| 0.375678 | 0.08981 |
| 0.377671 | 0.089219 |
| 0.379664 | 0.047391 |
| 0.381657 | 0.014645 |
| 0.38365 | 0.008371 |
| 0.385643 | -0.006932 |
| 0.387636 | 0.002422 |
| 0.389629 | -0.022229 |
| 0.391621 | -0.022466 |
| 0.393614 | -0.023651 |
| 0.395607 | -0.031165 |
| 0.3976 | 0.013307 |

|  | frequency |
| --- | --- |
| 0.000993451 | 1 |
| 0.00298035 | 1 |
| 0.00496726 | 1 |
| 0.00695416 | 1 |
| 0.00894106 | 1 |
| 0.010928 | 1 |
| 0.0129149 | 0.999994 |
| 0.0149018 | 0.999999 |
| 0.0168887 | 0.999989 |
| 0.0188756 | 0.999993 |
| 0.0208625 | 0.999978 |
| 0.0228494 | 0.999974 |
| 0.0248363 | 0.999952 |
| 0.0268232 | 0.99996 |
| 0.0288101 | 0.999944 |
| 0.030797 | 0.999963 |
| 0.0327839 | 0.999937 |
| 0.0347708 | 0.999938 |
| 0.0367577 | 0.999937 |
| 0.0387446 | 0.999842 |
| 0.0407315 | 0.999889 |
| 0.0427184 | 0.999889 |
| 0.0447053 | 0.99987 |
| 0.0466922 | 0.999821 |
| 0.0486791 | 0.999816 |
| 0.050666 | 0.999654 |
| 0.0526529 | 0.999593 |
| 0.0546398 | 0.999458 |
| 0.0566267 | 0.999362 |
| 0.0586136 | 0.999561 |
| 0.0606005 | 0.999531 |
| 0.0625874 | 0.999444 |
| 0.0645743 | 0.99949 |
| 0.0665612 | 0.999417 |
| 0.0685481 | 0.999411 |
| 0.070535 | 0.999395 |
| 0.0725219 | 0.999393 |
| 0.0745088 | 0.99927 |
| 0.0764957 | 0.999253 |
| 0.0784826 | 0.999411 |
| 0.0804695 | 0.999342 |
| 0.0824564 | 0.999546 |
| 0.0844433 | 0.99948 |
| 0.0864303 | 0.999554 |
| 0.0884172 | 0.999396 |
| 0.0904041 | 0.999465 |
| 0.092391 | 0.999389 |
| 0.0943779 | 0.999375 |
| 0.0963648 | 0.999373 |
| 0.0983517 | 0.999455 |
| 0.100339 | 0.999482 |
| 0.102325 | 0.999412 |
| 0.104312 | 0.999512 |
| 0.106299 | 0.99943 |
| 0.108286 | 0.99933 |
| 0.110273 | 0.999143 |
| 0.11226 | 0.999122 |
| 0.114247 | 0.998999 |
| 0.116234 | 0.998864 |
| 0.118221 | 0.998737 |
| 0.120208 | 0.998905 |
| 0.122194 | 0.998579 |
| 0.124181 | 0.998397 |
| 0.126168 | 0.998564 |
| 0.128155 | 0.998721 |
| 0.130142 | 0.99831 |
| 0.132129 | 0.998141 |
| 0.134116 | 0.997711 |
| 0.136103 | 0.997408 |
| 0.13809 | 0.997305 |
| 0.140077 | 0.997255 |
| 0.142064 | 0.997296 |
| 0.14405 | 0.997137 |
| 0.146037 | 0.997496 |
| 0.148024 | 0.997505 |
| 0.150011 | 0.997844 |
| 0.151998 | 0.99774 |
| 0.153985 | 0.997638 |
| 0.155972 | 0.997912 |
| 0.157959 | 0.997544 |
| 0.159946 | 0.997229 |
| 0.161933 | 0.997349 |
| 0.163919 | 0.99727 |
| 0.165906 | 0.997665 |
| 0.167893 | 0.997683 |
| 0.16988 | 0.997725 |
| 0.171867 | 0.997575 |
| 0.173854 | 0.997595 |
| 0.175841 | 0.997349 |
| 0.177828 | 0.99728 |
| 0.179815 | 0.997684 |
| 0.181802 | 0.997135 |
| 0.183788 | 0.997484 |
| 0.185775 | 0.997294 |
| 0.187762 | 0.997647 |
| 0.189749 | 0.997373 |
| 0.191736 | 0.996978 |
| 0.193723 | 0.997242 |
| 0.19571 | 0.996977 |
| 0.197697 | 0.997124 |
| 0.199684 | 0.997564 |
| 0.201671 | 0.997594 |
| 0.203657 | 0.997727 |
| 0.205644 | 0.997603 |
| 0.207631 | 0.997498 |
| 0.209618 | 0.997557 |
| 0.211605 | 0.997311 |
| 0.213592 | 0.997427 |
| 0.215579 | 0.997656 |
| 0.217566 | 0.99749 |
| 0.219553 | 0.99726 |
| 0.22154 | 0.996825 |
| 0.223527 | 0.996671 |
| 0.225513 | 0.996828 |
| 0.2275 | 0.996785 |
| 0.229487 | 0.996666 |
| 0.231474 | 0.99649 |
| 0.233461 | 0.996299 |
| 0.235448 | 0.996012 |
| 0.237435 | 0.995403 |
| 0.239422 | 0.995384 |
| 0.241409 | 0.994686 |
| 0.243396 | 0.994543 |
| 0.245382 | 0.994229 |
| 0.247369 | 0.993409 |
| 0.249356 | 0.993871 |
| 0.251343 | 0.993534 |
| 0.25333 | 0.99243 |
| 0.255317 | 0.991578 |
| 0.257304 | 0.990644 |
| 0.259291 | 0.98981 |
| 0.261278 | 0.989529 |
| 0.263265 | 0.989832 |
| 0.265251 | 0.989265 |
| 0.267238 | 0.98876 |
| 0.269225 | 0.988951 |
| 0.271212 | 0.987777 |
| 0.273199 | 0.986092 |
| 0.275186 | 0.985543 |
| 0.277173 | 0.98536 |
| 0.27916 | 0.984059 |
| 0.281147 | 0.982927 |
| 0.283134 | 0.982311 |
| 0.28512 | 0.980713 |
| 0.287107 | 0.978648 |
| 0.289094 | 0.97821 |
| 0.291081 | 0.977242 |
| 0.293068 | 0.975628 |
| 0.295055 | 0.973963 |
| 0.297042 | 0.969055 |
| 0.299029 | 0.967718 |
| 0.301016 | 0.964762 |
| 0.303003 | 0.963257 |
| 0.30499 | 0.961265 |
| 0.306976 | 0.957831 |
| 0.308963 | 0.956834 |
| 0.31095 | 0.955547 |
| 0.312937 | 0.955734 |
| 0.314924 | 0.950597 |
| 0.316911 | 0.949942 |
| 0.318898 | 0.946147 |
| 0.320885 | 0.94252 |
| 0.322872 | 0.935685 |
| 0.324859 | 0.931478 |
| 0.326845 | 0.924701 |
| 0.328832 | 0.913745 |
| 0.330819 | 0.910125 |
| 0.332806 | 0.90695 |
| 0.334793 | 0.90016 |
| 0.33678 | 0.887899 |
| 0.338767 | 0.883132 |
| 0.340754 | 0.878736 |
| 0.342741 | 0.872367 |
| 0.344728 | 0.861394 |
| 0.346714 | 0.853483 |
| 0.348701 | 0.833398 |
| 0.350688 | 0.807068 |
| 0.352675 | 0.81066 |
| 0.354662 | 0.797079 |
| 0.356649 | 0.779229 |
| 0.358636 | 0.765931 |
| 0.360623 | 0.74156 |
| 0.36261 | 0.733995 |
| 0.364597 | 0.72607 |
| 0.366583 | 0.710468 |
| 0.36857 | 0.691579 |
| 0.370557 | 0.686019 |
| 0.372544 | 0.662174 |
| 0.374531 | 0.651345 |
| 0.376518 | 0.640384 |
| 0.378505 | 0.616041 |
| 0.380492 | 0.620298 |
| 0.382479 | 0.59657 |
| 0.384466 | 0.553289 |
| 0.386453 | 0.525207 |
| 0.388439 | 0.519656 |
| 0.390426 | 0.519991 |
| 0.392413 | 0.510184 |
| 0.3944 | 0.49643 |
| 0.396387 | 0.454266 |

|  | frequency |
| --- | --- |
| 0.000993057 | 1 |
| 0.00297917 | 1 |
| 0.00496528 | 1 |
| 0.0069514 | 1 |
| 0.00893751 | 1 |
| 0.0109236 | 1 |
| 0.0129097 | 0.999999 |
| 0.0148958 | 1 |
| 0.016882 | 0.999999 |
| 0.0188681 | 1 |
| 0.0208542 | 0.999998 |
| 0.0228403 | 1 |
| 0.0248264 | 0.999999 |
| 0.0268125 | 1 |
| 0.0287986 | 0.999999 |
| 0.0307848 | 0.999997 |
| 0.0327709 | 0.999997 |
| 0.034757 | 0.999997 |
| 0.0367431 | 1 |
| 0.0387292 | 1 |
| 0.0407153 | 1 |
| 0.0427014 | 0.999999 |
| 0.0446875 | 0.999996 |
| 0.0466737 | 0.999999 |
| 0.0486598 | 0.999996 |
| 0.0506459 | 0.999995 |
| 0.052632 | 0.999992 |
| 0.0546181 | 0.999995 |
| 0.0566042 | 1 |
| 0.0585903 | 0.999997 |
| 0.0605764 | 0.999996 |
| 0.0625626 | 0.99999 |
| 0.0645487 | 0.999989 |
| 0.0665348 | 0.999997 |
| 0.0685209 | 0.999986 |
| 0.070507 | 0.999987 |
| 0.0724931 | 0.999995 |
| 0.0744792 | 0.999984 |
| 0.0764654 | 0.999989 |
| 0.0784515 | 0.999989 |
| 0.0804376 | 0.999985 |
| 0.0824237 | 0.999991 |
| 0.0844098 | 0.999982 |
| 0.0863959 | 0.999999 |
| 0.088382 | 0.999982 |
| 0.0903681 | 0.99999 |
| 0.0923543 | 0.999979 |
| 0.0943404 | 0.999993 |
| 0.0963265 | 0.999975 |
| 0.0983126 | 0.999988 |
| 0.100299 | 0.999988 |
| 0.102285 | 0.999986 |
| 0.104271 | 1 |
| 0.106257 | 0.999979 |
| 0.108243 | 0.999985 |
| 0.110229 | 0.999978 |
| 0.112215 | 0.999976 |
| 0.114202 | 0.999981 |
| 0.116188 | 0.999958 |
| 0.118174 | 0.999956 |
| 0.12016 | 0.999961 |
| 0.122146 | 0.999961 |
| 0.124132 | 0.999965 |
| 0.126118 | 0.999962 |
| 0.128104 | 0.999965 |
| 0.13009 | 0.99995 |
| 0.132077 | 0.999947 |
| 0.134063 | 0.999959 |
| 0.136049 | 0.999954 |
| 0.138035 | 0.999948 |
| 0.140021 | 0.999933 |
| 0.142007 | 0.999958 |
| 0.143993 | 0.999958 |
| 0.145979 | 0.999958 |
| 0.147965 | 0.99994 |
| 0.149952 | 0.999916 |
| 0.151938 | 0.999947 |
| 0.153924 | 0.999952 |
| 0.15591 | 0.99993 |
| 0.157896 | 0.99996 |
| 0.159882 | 0.999943 |
| 0.161868 | 0.999939 |
| 0.163854 | 0.999926 |
| 0.16584 | 0.999929 |
| 0.167827 | 0.999931 |
| 0.169813 | 0.999941 |
| 0.171799 | 0.999953 |
| 0.173785 | 0.99994 |
| 0.175771 | 0.999933 |
| 0.177757 | 0.999939 |
| 0.179743 | 0.999938 |
| 0.181729 | 0.999939 |
| 0.183715 | 0.999927 |
| 0.185702 | 0.999928 |
| 0.187688 | 0.999939 |
| 0.189674 | 0.999934 |
| 0.19166 | 0.999944 |
| 0.193646 | 0.999935 |
| 0.195632 | 0.999934 |
| 0.197618 | 0.999929 |
| 0.199604 | 0.99994 |
| 0.20159 | 0.999944 |
| 0.203577 | 0.999942 |
| 0.205563 | 0.999925 |
| 0.207549 | 0.999966 |
| 0.209535 | 0.999946 |
| 0.211521 | 0.999942 |
| 0.213507 | 0.999922 |
| 0.215493 | 0.999923 |
| 0.217479 | 0.999945 |
| 0.219465 | 0.99994 |
| 0.221452 | 0.999945 |
| 0.223438 | 0.999918 |
| 0.225424 | 0.999925 |
| 0.22741 | 0.999932 |
| 0.229396 | 0.999931 |
| 0.231382 | 0.999912 |
| 0.233368 | 0.999921 |
| 0.235354 | 0.999936 |
| 0.237341 | 0.999926 |
| 0.239327 | 0.999919 |
| 0.241313 | 0.999903 |
| 0.243299 | 0.999884 |
| 0.245285 | 0.999867 |
| 0.247271 | 0.999885 |
| 0.249257 | 0.999887 |
| 0.251243 | 0.999799 |
| 0.253229 | 0.999822 |
| 0.255216 | 0.999836 |
| 0.257202 | 0.999807 |
| 0.259188 | 0.999766 |
| 0.261174 | 0.999761 |
| 0.26316 | 0.999772 |
| 0.265146 | 0.999731 |
| 0.267132 | 0.999735 |
| 0.269118 | 0.999713 |
| 0.271104 | 0.999703 |
| 0.273091 | 0.99963 |
| 0.275077 | 0.999641 |
| 0.277063 | 0.999594 |
| 0.279049 | 0.999656 |
| 0.281035 | 0.999586 |
| 0.283021 | 0.999516 |
| 0.285007 | 0.999487 |
| 0.286993 | 0.999427 |
| 0.288979 | 0.999378 |
| 0.290966 | 0.99938 |
| 0.292952 | 0.999415 |
| 0.294938 | 0.999358 |
| 0.296924 | 0.999277 |
| 0.29891 | 0.999305 |
| 0.300896 | 0.999194 |
| 0.302882 | 0.999103 |
| 0.304868 | 0.999065 |
| 0.306854 | 0.998913 |
| 0.308841 | 0.998923 |
| 0.310827 | 0.998755 |
| 0.312813 | 0.998635 |
| 0.314799 | 0.99861 |
| 0.316785 | 0.998566 |
| 0.318771 | 0.998444 |
| 0.320757 | 0.998336 |
| 0.322743 | 0.998241 |
| 0.324729 | 0.9982 |
| 0.326716 | 0.998129 |
| 0.328702 | 0.997747 |
| 0.330688 | 0.997588 |
| 0.332674 | 0.997345 |
| 0.33466 | 0.997102 |
| 0.336646 | 0.99692 |
| 0.338632 | 0.996781 |
| 0.340618 | 0.996623 |
| 0.342605 | 0.996217 |
| 0.344591 | 0.996134 |
| 0.346577 | 0.995823 |
| 0.348563 | 0.995362 |
| 0.350549 | 0.994573 |
| 0.352535 | 0.994271 |
| 0.354521 | 0.993728 |
| 0.356507 | 0.992665 |
| 0.358493 | 0.992227 |
| 0.36048 | 0.991455 |
| 0.362466 | 0.990854 |
| 0.364452 | 0.990614 |
| 0.366438 | 0.989317 |
| 0.368424 | 0.988426 |
| 0.37041 | 0.987565 |
| 0.372396 | 0.987639 |
| 0.374382 | 0.986349 |
| 0.376368 | 0.985591 |
| 0.378355 | 0.984157 |
| 0.380341 | 0.983459 |
| 0.382327 | 0.981333 |
| 0.384313 | 0.979068 |
| 0.386299 | 0.977685 |
| 0.388285 | 0.976136 |
| 0.390271 | 0.973545 |
| 0.392257 | 0.971437 |
| 0.394243 | 0.970084 |
| 0.39623 | 0.968194 |

|  | frequency |
| --- | --- |
| 0.000993451 | 1 |
| 0.00298035 | 1 |
| 0.00496726 | 1 |
| 0.00695416 | 1 |
| 0.00894106 | 1 |
| 0.010928 | 1 |
| 0.0129149 | 1 |
| 0.0149018 | 0.999999 |
| 0.0168887 | 1 |
| 0.0188756 | 1 |
| 0.0208625 | 0.999998 |
| 0.0228494 | 1 |
| 0.0248363 | 0.999998 |
| 0.0268232 | 0.999998 |
| 0.0288101 | 0.999997 |
| 0.030797 | 0.999993 |
| 0.0327839 | 0.999996 |
| 0.0347708 | 1 |
| 0.0367577 | 0.999998 |
| 0.0387446 | 0.999996 |
| 0.0407315 | 0.99999 |
| 0.0427184 | 0.999996 |
| 0.0447053 | 0.999991 |
| 0.0466922 | 0.999993 |
| 0.0486791 | 0.999991 |
| 0.050666 | 0.999984 |
| 0.0526529 | 0.999981 |
| 0.0546398 | 0.999972 |
| 0.0566267 | 0.99998 |
| 0.0586136 | 0.999985 |
| 0.0606005 | 0.99998 |
| 0.0625874 | 0.999971 |
| 0.0645743 | 0.999967 |
| 0.0665612 | 0.999979 |
| 0.0685481 | 0.999971 |
| 0.070535 | 0.999974 |
| 0.0725219 | 0.999974 |
| 0.0745088 | 0.999977 |
| 0.0764957 | 0.999968 |
| 0.0784826 | 0.999979 |
| 0.0804695 | 0.999963 |
| 0.0824564 | 0.99997 |
| 0.0844433 | 0.999974 |
| 0.0864303 | 0.999967 |
| 0.0884172 | 0.999967 |
| 0.0904041 | 0.999969 |
| 0.092391 | 0.999973 |
| 0.0943779 | 0.999976 |
| 0.0963648 | 0.999981 |
| 0.0983517 | 0.999977 |
| 0.100339 | 0.999978 |
| 0.102325 | 0.999971 |
| 0.104312 | 0.999978 |
| 0.106299 | 0.999992 |
| 0.108286 | 0.999971 |
| 0.110273 | 0.999975 |
| 0.11226 | 0.999979 |
| 0.114247 | 0.999961 |
| 0.116234 | 0.999939 |
| 0.118221 | 0.999957 |
| 0.120208 | 0.999971 |
| 0.122194 | 0.999963 |
| 0.124181 | 0.999939 |
| 0.126168 | 0.999946 |
| 0.128155 | 0.999936 |
| 0.130142 | 0.999934 |
| 0.132129 | 0.999934 |
| 0.134116 | 0.999925 |
| 0.136103 | 0.999927 |
| 0.13809 | 0.999933 |
| 0.140077 | 0.999944 |
| 0.142064 | 0.999916 |
| 0.14405 | 0.999931 |
| 0.146037 | 0.999922 |
| 0.148024 | 0.999912 |
| 0.150011 | 0.999917 |
| 0.151998 | 0.999926 |
| 0.153985 | 0.999915 |
| 0.155972 | 0.999926 |
| 0.157959 | 0.999925 |
| 0.159946 | 0.999914 |
| 0.161933 | 0.999888 |
| 0.163919 | 0.99992 |
| 0.165906 | 0.999913 |
| 0.167893 | 0.99992 |
| 0.16988 | 0.999925 |
| 0.171867 | 0.999918 |
| 0.173854 | 0.999915 |
| 0.175841 | 0.999919 |
| 0.177828 | 0.999878 |
| 0.179815 | 0.999919 |
| 0.181802 | 0.999914 |
| 0.183788 | 0.999911 |
| 0.185775 | 0.999911 |
| 0.187762 | 0.999892 |
| 0.189749 | 0.999917 |
| 0.191736 | 0.999913 |
| 0.193723 | 0.999917 |
| 0.19571 | 0.999921 |
| 0.197697 | 0.999917 |
| 0.199684 | 0.999912 |
| 0.201671 | 0.999899 |
| 0.203657 | 0.999918 |
| 0.205644 | 0.999933 |
| 0.207631 | 0.999936 |
| 0.209618 | 0.999933 |
| 0.211605 | 0.999904 |
| 0.213592 | 0.99992 |
| 0.215579 | 0.999917 |
| 0.217566 | 0.999895 |
| 0.219553 | 0.999898 |
| 0.22154 | 0.999894 |
| 0.223527 | 0.999896 |
| 0.225513 | 0.999917 |
| 0.2275 | 0.999878 |
| 0.229487 | 0.999884 |
| 0.231474 | 0.999888 |
| 0.233461 | 0.999894 |
| 0.235448 | 0.999891 |
| 0.237435 | 0.999896 |
| 0.239422 | 0.999884 |
| 0.241409 | 0.999866 |
| 0.243396 | 0.999836 |
| 0.245382 | 0.999815 |
| 0.247369 | 0.999789 |
| 0.249356 | 0.999805 |
| 0.251343 | 0.999779 |
| 0.25333 | 0.999762 |
| 0.255317 | 0.999731 |
| 0.257304 | 0.999719 |
| 0.259291 | 0.999677 |
| 0.261278 | 0.999672 |
| 0.263265 | 0.999713 |
| 0.265251 | 0.999634 |
| 0.267238 | 0.999637 |
| 0.269225 | 0.99962 |
| 0.271212 | 0.999642 |
| 0.273199 | 0.999605 |
| 0.275186 | 0.999576 |
| 0.277173 | 0.99954 |
| 0.27916 | 0.999477 |
| 0.281147 | 0.999443 |
| 0.283134 | 0.999374 |
| 0.28512 | 0.999376 |
| 0.287107 | 0.999292 |
| 0.289094 | 0.999278 |
| 0.291081 | 0.99923 |
| 0.293068 | 0.999211 |
| 0.295055 | 0.999189 |
| 0.297042 | 0.999057 |
| 0.299029 | 0.998969 |
| 0.301016 | 0.998913 |
| 0.303003 | 0.998833 |
| 0.30499 | 0.998723 |
| 0.306976 | 0.998625 |
| 0.308963 | 0.998616 |
| 0.31095 | 0.998535 |
| 0.312937 | 0.998479 |
| 0.314924 | 0.998371 |
| 0.316911 | 0.998369 |
| 0.318898 | 0.998216 |
| 0.320885 | 0.998058 |
| 0.322872 | 0.997822 |
| 0.324859 | 0.997698 |
| 0.326845 | 0.997463 |
| 0.328832 | 0.997122 |
| 0.330819 | 0.99694 |
| 0.332806 | 0.996643 |
| 0.334793 | 0.996476 |
| 0.33678 | 0.995939 |
| 0.338767 | 0.995564 |
| 0.340754 | 0.995269 |
| 0.342741 | 0.995264 |
| 0.344728 | 0.994585 |
| 0.346714 | 0.994239 |
| 0.348701 | 0.993719 |
| 0.350688 | 0.992898 |
| 0.352675 | 0.992289 |
| 0.354662 | 0.991558 |
| 0.356649 | 0.990998 |
| 0.358636 | 0.990263 |
| 0.360623 | 0.988535 |
| 0.36261 | 0.988274 |
| 0.364597 | 0.987892 |
| 0.366583 | 0.987182 |
| 0.36857 | 0.986203 |
| 0.370557 | 0.985163 |
| 0.372544 | 0.983986 |
| 0.374531 | 0.98446 |
| 0.376518 | 0.983115 |
| 0.378505 | 0.980343 |
| 0.380492 | 0.977934 |
| 0.382479 | 0.975921 |
| 0.384466 | 0.973359 |
| 0.386453 | 0.971219 |
| 0.388439 | 0.970332 |
| 0.390426 | 0.968077 |
| 0.392413 | 0.964392 |
| 0.3944 | 0.961012 |
| 0.396387 | 0.95931 |

EV-D68 MO native virion

EV-D68 MO in complex with inhibitor 11526092

EV-D68 MO in complex with pleconaril

EV-D68 MO in complex with inhibitor 11526091

EV-D68 MO in complex with inhibitor 11526093

**Tables**

**Table S1.** Calculated molecule properties for pleconaril and analogs using ChemAxon software (Budapest, Hungary). PSA = polar surface area, HBA = hydrogen bond acceptors, HBD = hydrogen bond donors, MW = molecular weight, MPO = multiparameter optimization (BBB score).

| Compound | PSA | HBA | HBD | logP | logD  [pH=7.4] | MW | pKa | MPO | MPOv2 |
| --- | --- | --- | --- | --- | --- | --- | --- | --- | --- |
| 11526091 | 137.63 | 7 | 0 | 3.43 | 3.43 | 455.35 | -1.95 | 3.39 | 4.10 |
| 11526092 | 94.49 | 5 | 0 | 3.97 | 3.97 | 424.37 | -1.93 | 3.92 | 4.90 |
| 11526093 | 94.49 | 5 | 0 | 3.62 | 3.62 | 428.34 | -1.94 | 4.24 | 5.05 |
| Pleconaril | 74.18 | 4 | 0 | 5.04 | 5.04 | 381.35 | 1.64 | 3.85 | 4.85 |

**Table S2.** Results for maximum tolerated dose studies.

| **Age** | **Dose (mg/kg/day)** | **No. of animals** | **Survival** | **Difference between survival?** | **Treatment Schedule** | **Treatment Route** | **% Weight Gain vs. Vehicle** |
| --- | --- | --- | --- | --- | --- | --- | --- |
| 4-week-old | 10 | 5 | 5/5 | NS | b.i.d. x 5 days | PO | NS |
|  | 30 |  | 5/5 | NS |  |  | NS |
|  | 100 |  | 5/5 | NS |  |  | NS |
| 10-day-old | 1 |  | 5/5 | NS |  | IP | Yes^*^ |
|  | 3 |  | 4/5 | NS |  |  | Yes^*^ |
|  | 10 |  | 5/5 | NS |  |  | NS |

**Table S3.** Experimental Design (AFM respiratory model).

| **Number/Cage** | **Infected** | **Compound** | **Dosage** | **Route** | **Treatment Schedule** | **Observations** |
| --- | --- | --- | --- | --- | --- | --- |
| 12 | Yes | Placebo* | - | PO | q.d. x 5 days beginning 2 hr pre-infection | 4 mice per group sacrificed at days 1, 3, and 5 post infection  for: lung virus titers and cytokines, and blood virus titers |
|  |  | 11526092 | 10 mg/kg/d |  |  |  |
|  |  |  | 30 mg/kg/d |  |  |  |
|  |  |  | 100 mg/kg/d |  |  |  |
|  |  | Guanidine | 100 mg/kg/d | IP | b.i.d. x 5 days beginning 4 hr post-infection |  |
| 5 | No | Normal Controls | | Untreated mice observed for normal weight gain. | | |

**Table S4.** Experimental Design (AFM neurological models 4h post infection).

| **Number/Cage** | **Infected** | **Compound** | **Dosage** | **Route** | **Treatment Schedule** | **Observations** |
| --- | --- | --- | --- | --- | --- | --- |
| 8 | Yes | Placebo* | - | IP | b.i.d. x 5 days beginning 4 hr post-infection | Mice observed for mortality, weight loss and neurological scores for 21 days.  Blood collected from 3^**^ mice/group on days 1,3 and 5 from determination of viremia |
|  |  | 11526092 | 1 mg/kg/d |  |  |  |
|  |  |  | 3 mg/kg/d |  |  |  |
|  |  |  | 10 mg/kg/d |  |  |  |
|  |  | Guanidine | 100 mg/kg/d |  |  |  |
| 2 | No | Normal Controls | | Untreated mice observed for normal weight gain. | | |
| *A suspension of 0.5% carboxymethylcellulose (CMC) and 0.5% Tween 20 in sterile water was used as the vehicle  **Blood collection for viremia from 3 mice/day-different mice to be bled on successive days due to young age | | | | | | |

**Table S5.** Experimental Design (AFM neurological model pre-exposure treatment).

| **Number** | **Infected** | **Compound** | **Dosage** | **Route** | **Treatment Schedule** | **Observations** |
| --- | --- | --- | --- | --- | --- | --- |
| 8 | Yes | Placebo* | - | IP | b.i.d. x 5 days beginning 2 hr pre-infection | Mice observed for mortality, weight loss and neurological scores for 21 days.  Blood collected from 3^**^ mice/group on days 1,3 and 5 from determination of viremia |
|  |  | 11526092 | 1 mg/kg/d |  |  |  |
|  |  |  | 3 mg/kg/d |  |  |  |
|  |  |  | 10 mg/kg/d |  |  |  |
|  |  | Guanidine | 100 mg/kg/d |  |  |  |
| 2 | No | Normal Controls | | Untreated mice observed for normal weight gain. | | |
| *A suspension of 0.5% carboxymethylcellulose (CMC) and 0.5% Tween 20 in sterile water was used as the vehicle  **Blood collection for viremia from 3 mice/day-different mice to be bled on successive days due to young age | | | | | | |

**Table S6.** Validation for cryo-EM.

|  | **EV-D68 MO control** | **EV-D68 MO and**  **11526092** | **EV-D68 MO and**  **Pleconaril** |
| --- | --- | --- | --- |
| **Cryo-EM data acquisition and processing** | | |  |
| Magnification | 64,000X | 64,000X | 64,000X |
| Camera | K3 Direct Detection Camera (Gatan) | K3 Direct Detection Camera (Gatan) | K3 Direct Detection Camera (Gatan) |
| Voltage (kV) | 300 | 300 | 300 |
| Pixel size (Å) | 0.666 | 0.666 | 0.664 |
| Defocus range (µm) | 0.4-2 | 0.4-2 | 0.4-2 |
| Total electron dose (electrons/Å^2^ ) | 36.06 | 36.06 | 36.3 |
| Particles picked | 360,944 | 963,036 | 355,591 |
| Particles used | 239,843 | 452,877 | 355,591 |
| Map resolution  (FSC threshold =0.143) | 2 | 2 | 2.7 |
| **Model building and refinement** | | |  |
| Model building reference  (PDB code) | 6CSG | 6CSG | 6CSG |
| MolProbity score | 1.24 | 1.35 | 1.95 |
| Clashscore | 3.57 | 5.62 | 13.58 |
| Rotamer outliers (%) | 1.00 | 1.14 | 0.00 |
| R.m.s. deviations | | |  |
| Bond length (Å) | 0.002 | 0.002 | 0.003 |
| Bond angles (°) | 0.540 | 0.542 | 0.534 |
| Ramachandran plot (%) | | |  |
| Favored | 97.54 | 98.15 | 95.52 |
| Allowed | 2.46 | 1.85 | 4.48 |
| Outliers | 0.00 | 0.00 | 0.00 |

|  | **EV-D68 MO and**  **11526091** | **EV-D68 MO and**  **11526093** |
| --- | --- | --- |
| **Cryo-EM data acquisition and processing** | | |
| Magnification | 64,000X | 64,000X |
| Camera | K3 Direct Detection Camera (Gatan) | K3 Direct Detection Camera (Gatan) |
| Voltage (kV) | 300 | 300 |
| Pixel size (Å) | 0.666 | 0.666 |
| Defocus range (µm) | 0.4-2 | 0.4-2 |
| Total electron dose (electrons/Å^2^ ) | 36.06 | 36.06 |
| Particles picked | 22,634 | 604,231 |
| Particles used | 20,971 | 225,145 |
| Map resolution  (FSC threshold =0.143) | 2.3 | 2 |
| **Model building and refinement** | | |
| Model building reference  (PDB code) | 6CSG | 6CSG |
| MolProbity score | 1.57 | 1.40 |
| Clashscore | 4.44 | 2.85 |
| Rotamer outliers (%) | 2.84 | 1.71 |
| R.m.s. deviations | | |
| Bond length (Å) | 0.002 | 0.002 |
| Bond angles (°) | 0.511 | 0.560 |
| Ramachandran plot (%) | | |
| Favored | 98.28 | 97.17 |
| Allowed | 1.72 | 2.83 |
| Outliers | 0.00 | 0.00 |
